# CoSMoMVPA: multi-modal multivariate pattern analysis of neuroimaging data in Matlab / GNU Octave

**DOI:** 10.1101/047118

**Authors:** Nikolaas N. Oosterhof, Andrew C. Connolly, James V. Haxby

**Keywords:** multi-variate pattern analysis, software, functional magnetic resonance imaging, magnetoencephalography, electroencephalography, open source, cognitive neuroscience

## Abstract

Recent years have seen an increase in the popularity of multivariate pattern (MVP) analysis of functional magnetic resonance (fMRI) data, and, to a much lesser extent, magneto-and electro-encephalography (M/EEG) data. We present CoSMoMVPA, a lightweight MVPA (MVP analysis) toolbox implemented in the intersection of the Matlab and GNU Octave languages, that treats both fMRI and M/EEG data as first-class citizens.

CoSMoMVPA supports all state-of-the-art MVP analysis techniques, including searchlight analyses, classification, correlations, representational similarity analysis, and the time generalization method. These can be used to address both data-driven and hypothesis-driven questions about neural organization and representations, both within and across: space, time, frequency bands, neuroimaging modalities, individuals, and species.

It uses a uniform data representation of fMRI data in the volume or on the surface, and of M/EEG data at the sensor and source level. Through various external toolboxes, it directly supports reading and writing a variety of fMRI and M/EEG neuroimaging formats, and, where applicable, can convert between them. As a result, it can be integrated readily in existing pipelines and used with existing preprocessed datasets.

CoSMoMVPA overloads the traditional volumetric searchlight concept to support neighborhoods for M/EEG and surface-based fMRI data, which supports localization of multivariate effects of interest across space, time, and frequency dimensions. CoSMoMVPA also provides a generalized approach to multiple comparison correction across these dimensions using Threshold-Free Cluster Enhancement with state-of-the-art clustering and permutation techniques.

CoSMoMVPA is highly modular and uses abstractions to provide a uniform interface for a variety of MVP measures. Typical analyses require a few lines of code, making it accessible to beginner users. At the same time, expert programmers can easily extend its functionality.

CoSMoMVPA comes with extensive documentation, including a variety of runnable demonstration scripts and analysis exercises (with example data and solutions). It uses best software engineering practices including version control, distributed development, an automated test suite, and continuous integration testing. It can be used with the proprietary Matlab and the free GNU Octave software, and it complies with open source distribution platforms such as NeuroDebian.

CoSMoMVPA is Free/Open Source Software under the permissive MIT license.

Website: https://cosmomvpa.org

Source code: https://github.com/CoSMoMVPA/CoSMoMVPA

## 1 Introduction

The use of multivariate pattern analysis has in the last decade become popular in functional magnetic resonance imaging (fMRI) research (Edelman, Grill-Spector, Kushnir, & Malach, 1998; Haxby et al., 2001; D. D. Cox & Savoy, 2003; Mitchell et al., 2004; Haynes & Rees, 2006; Norman, Polyn, Detre, & Haxby, 2006). This is not surprising, as MVPA has several advantages compared to traditional, and more commonly used, univariate analyses. First, MVPA can provide more sensitivity in discriminating conditions of interest than univariate approaches because it considers patterns of voxel activity that may show weak but consistent differences between conditions. Second, it allows for making inferences about the underlying neural representations, within (Peelen, Wiggett, & Downing, 2006) and across (Haxby et al., 2011) individuals, imaging modalities, and species (Kriegeskorte, Mur, Ruff, et al., 2008; Kriegeskorte & Kievit, 2013). The popularity of MVPA has resulted in several projects, such as PyMVPA (Hanke, Halchenko, Sederberg, Hanson, et al., 2009; Hanke, Halchenko, Sederberg, Olivetti, et al., 2009), the Princeton MVPA toolbox (Detre et al., 2006), PRoNTO (Schrouff et al., 2013), Searchmight (Pereira & Botvinick, 2011), the RSA toolbox (Nili et al., 2014) and The Decoding Toolbox (Hebart, Görgen, & Haynes, 2014), that aim to provide a common framework to make it accessible to non-expert programmers.

While MVPA has become a popular technique for fMRI, there are few reports of its application to magneto-and electro-encephalography (M/EEG ^1^) data (but see for example Kauhanen et al., 2006, Perreau Guimaraes, Wong, Uy, Grosenick, & Suppes, 2007, Pistohl, Schulze-Bonhage, & Aertsen, 2011, Chan, Halgren, Marinkovic, & Cash, 2011, Cichy, Pantazis, & Oliva, 2014). M/EEG data benefits from of a much higher temporal resolution than fMRI data, which allows for identifying neural correlates over the temporal dimension with high precision. In particular the time-generalization method (King & Dehaene, 2014) provides a promising avenue to investigate how different neural populations may encode experimental conditions over time. At the same time, to our knowledge only the MNE-python package (Gramfort et al., 2013) provides (limited) MVPA support for M/EEG data.

To address this gap, we present CoSMoMVPA, a toolbox that can be used to answer hypothesis-driven and data-driven neuroimaging questions using MVPA applied to both fMRI and M/EEG data. It uses simple, yet powerful data structures and a modular approach, so that different modules can be combined to build complex analysis pipelines. For data-driven approaches, it provides a generalized searchlight for localization of effects across voxels, surface nodes, M/EEG channels, time intervals, and frequency intervals; and combinations of those. To guard against type-1 errors, it it supports state-of-the-art Threshold-Free Cluster Enhancement and Monte Carlo-based permutation testing to correct for multiple comparisons.

It has been proposed (Hanke, Halchenko, Sederberg, Hanson, et al., 2009) that a framework for neuroimaging analysis should have at least the following five features:

- *intuitive user interface*: because most cognitive neuroscientists have limited training as computer scientists, the software should support workflows for common data analysis pipelines.
- *extensibility*: to avoid duplication of implementation efforts, the software should be able to interface with existing toolboxes.
- *transparent I/O*: the software should support easy input/output from and to neuroimaging data stored in common formats.
- *portability*: the software should not impose restrictions on the type of hardware used, and should run on all major operating systems.
- *open source software*: the software itself should be open to inspection of the implementation so that its correctness can be verified. In addition, we propose that the software should also run on open source platforms, so that researchers are not limited by the lack of access to proprietary software.

Yet another MVPA toolbox? CoSMoMVPA is not the first MVPA toolbox for cognitive neuroscience; indeed, since the Princeton toolbox (Detre et al., 2006), several other toolboxes have been released, both for the Python lan-guage—PyMVPA (Hanke, Halchenko, Sederberg, Hanson, et al., 2009; Hanke, Halchenko, Sederberg, Olivetti, et al., 2009), MNE-python (Gramfort et al., 2013)—and the Matlab language—PRoNTO (Schrouff et al., 2013), Search-might (Pereira & Botvinick, 2011), the RSA toolbox (Nili et al., 2014) and The Decoding Toolbox (Hebart et al., 2014). However, we think that CoSMoMVPA provides a unique combination of features that make it attractive for cognitive neuroscientists.

- *easy to use*: CoSMoMVPA is based on a small number of concepts (for details see Section 3), which apply directly state-of-the art MVP analyses using only a few lines of code, as illustrated in Section 2. This includes classification, representational similarity, and classification analyses; using either a region-of-interest or a searchlight; for fMRI and/or M/EEG data. Particular attention has been given to extensive data validation and, whereever necessary, providing informative error messages.
- *easy to learn*: CoSMoMVPA’s website provides a set of exercises (with solutions) to get familiar with CoSMoMVPA’s concepts. CoSMoMVPA also comes with a variety of runnable code examples for MVP analyses, a subset of which is shown in Section 2; users can use these as a starting point and adapt these examples for their own analyses.
- *characterize ‘locations’ that carry information*: CoSMOMVPA provides functionality for characterizing information content through a general searchlight concept. A searchlight is used with neighborhoods defined over space—voxels for volumetric fMRI or M/EEG source data, nodes for surface-based fMRI data, or channels for M/EEG channel data)—and time—timepoints for fMRI and M/EEG data, and temporal oscillations (frequency bands) for M/EEG data—to characterize information content over space and time.
- *based on Matlab / GNU Octave*: Matlab (and to lesser extent, GNU Octave) is a popular platform in cognitive neuroscience research, with many other widely used packages running on it, including Psychtoolbox (Brainard, 1997), FieldTrip (Oostenveld, Fries, Maris, & Schoffelen, 2011), EEGLAB (Delorme & Makeig, 2004), and SPM (Friston, Jezzard, & Turner, 1994). It is also a popular tool for general data analysis. Given its widespread use, many researchers may already be familiar with the Matlab / GNU Octave language, resulting in a reduced time-investment to learn CoSMoMVPA.
- *integrates with other packages*: CoSMoMVPA provides extensive input/output support. It can read and write the general fMRI NIFTI and GIFTI formats, and in addition read and write in the native data formats used by the AFNI (R. W. Cox, 1996), BrainVoyager (Goebel, Esposito, & Formisano, 2006), SPM (Friston et al., 1994), FieldTrip (Oostenveld et al., 2011) and EEGLAB (Delorme & Makeig, 2004) packages. As a result, it can be easily integrated in, or extend, existing data analysis pipelines.
- *focus on reproducibility and maintainability*: Inspired by PyMVPA, CoSMoMVPA uses the git distributed version control system (Torvalds et al., 2005)), an extenstive test suite, and continuous integration testing. These components improve maintainibility of the software, as improvements of the code can be made in a distributed manner, changes can be tracked over time, and (because of automated and repeated testing) changes that break existing functionality is likely to be detected very early by the developers. Since CoSMoMVPA runs on Open Source software, all components, at any point in their lifetime, can be studied and their behaviour reproduced in arbitrary detail (For details, see Section 4).

The remainder of this paper is as follows. Section 2 contains a series of motivating examples of analysis of fMRI and M/EEG data. Section 3 explains in more detail the CoSMoMVPA concepts underlying these examples. Section 4 explains some design decisions. Section 5 concludes the paper.

## 2 Analysis examples

This section provides a series of motivating examples of CoSMoMVPA’s approach to MVPA. To anticipate section 3, the examples use a variety of CoSMoMVPA concepts, including measures, neighborhoods, and searchlights. These examples demonstrate common MVP analyses, such as classification, correlation, representational similarity analysis, and the time generalization method. The examples are minor variations of the examples that are included with CoSMoMVPA, and based on real fMRI and M/EEG data. All data used here in the analyses were measured from participants who gave informed consent for procedures approved by the Ethical Committee of the University of Trento and/or the Institutional Review Board at Dartmouth College. The data is provided under a permissive license from the CoSMoMVPA website.

### 2.1 Classification

Data for our first example is from an fMRI experiment where one participant pressed the index and middle finger during different blocks. The experiment had four runs, each with four blocks for each finger. The data was preprocessed and analyzed with the general linear model in AFNI (R. W. Cox, 1996), resulting in t-statistics for each block. The ‘ingredients’ that the user has to specify are the filename of the AFNI neuroimaging data file, the `chunks`, which is the acquisition run number, and `targets`, which specifies which finger was pressed during each block. Classification analysis then requires specifying the cross-validation measure, classifier, and partitioning scheme. For surface-based analysis, the additional information needed consists of the surfaces used to delineate the grey matter and the number of voxels to select in each searchlight.

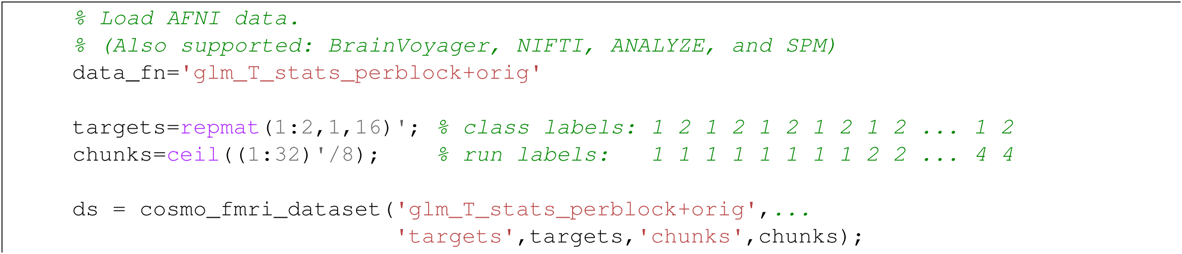

To run cross-validation using a classifier (e.g. D. D. Cox & Savoy, 2003), a cross-validation measure uses a take-one-run out cross-validation and an LDA classifier. Both the measure and the classifier are specified using a function handle; as illustrated in other examples (below), different measures or classifiers can be used by specifying another function handle.

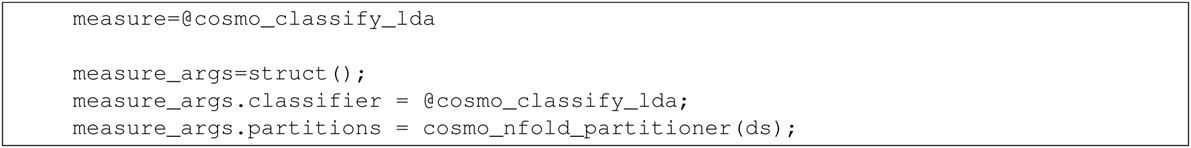

For ROI analysis in a user-defined mask, the whole-brain dataset is sliced using the mask resulting in a smaller dataset, and the measure applied to the smaller dataset.

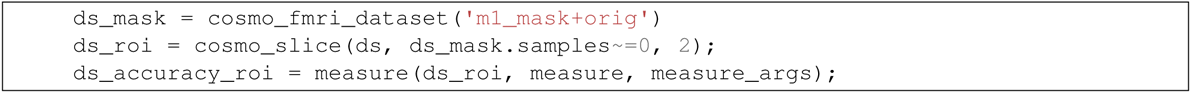

Surface-based searchlight analyses require one or two surfaces that define where the grey matter is. This example uses two surfaces from FreeSurfer that define the inner and outer grey-matter boundaries. Thus, each searchlight is shaped as a curved cylinder in between the two surfaces. Here, the number of voxels is kept constant across searchlight locations, therefore resulting in a variable radius across locations (due to variations in grey-matter thickness). A searchlight map (Kriegeskorte, Goebel, & Bandettini, 2006; Oosterhof, Wiestler, Downing, & Diedrichsen, 2011) is computed using `cosmo_searchlight`, which uses the neighborhood, measure, and the measure’s arguments. For illustration purposes, the result is stored in the GIFTI, AFNI / SUMA, and BrainVoyager formats.

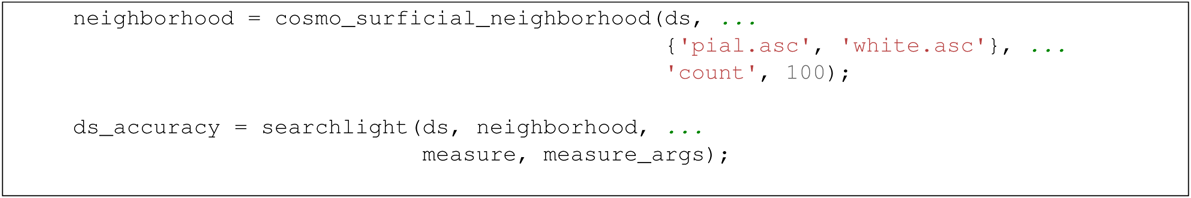

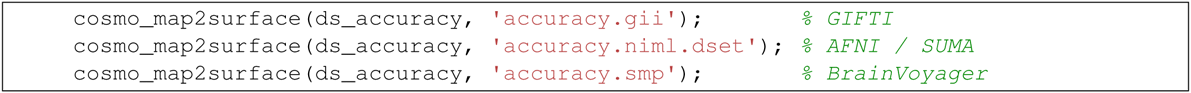

**Fig. 1:**
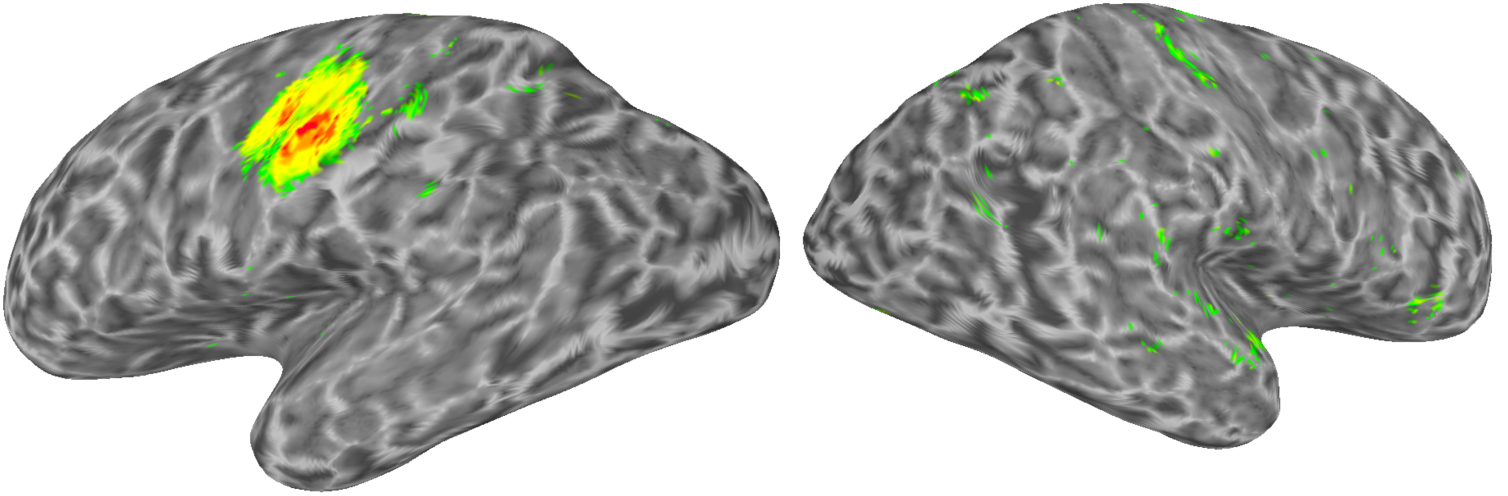
Illustration of surface-based searchlight results identifying where in the brain presses of the index and middle finger can be distinguished using an LDA classifier and take-one-run-out cross-validation.

Classification options can be set by adjusting the measure’s argument. For example, the following code defines using an odd-even partitioning scheme, SVM classifier, and output with the predicted label for each sample instead of classification accuracies:

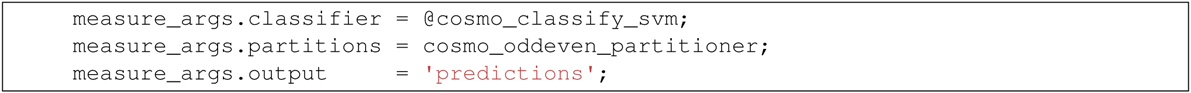

Region-of-interest and searchlight analyses with these parameters would then proceed with exactly the same calls to the measure and searchlight function as above.

### 2.2 Representational similarity analysis

Using data from Connolly et al., 2012, this example illustrates representational similarity analysis (Kriegeskorte, Mur, & Bandettini, 2008). The data has t-statistics from six categories (monkey, lemur, mallard, warbler, ladybug, lunamoth) averaged across all runs. Data can be loaded directly using `cosmo_fmri_dataset`, and a brain mask can be specified to only select voxels inside the brain. The only additional step is setting the targets (condition labels) to six unique values, and chunks to all the same value.

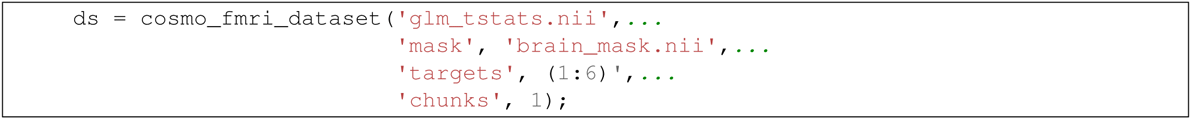

Behavioural data was collected where participants indicated the pair-wise similarity across categories, and stored in a matrix as illustrated here:

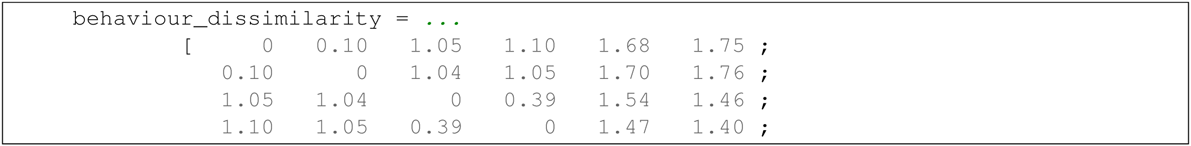

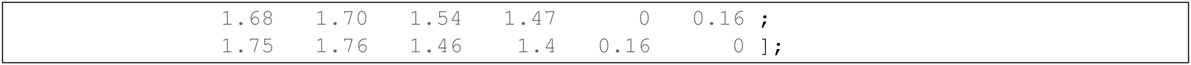

Using the behavioural dissimilarity matrix and a measure that computes similarity between the matrix and the neural dissimilarity, a searchlight can identify regions where the neural dissimilarity matches behavioural dissimilarity. The searchlight uses a spherical neighborhood with a variable radius, selecting the 100 voxels nearest to each center voxel. Using a fixed number of voxels ensures that the number of features for locations near the edge of the brain is the same as for locations inside the brain.

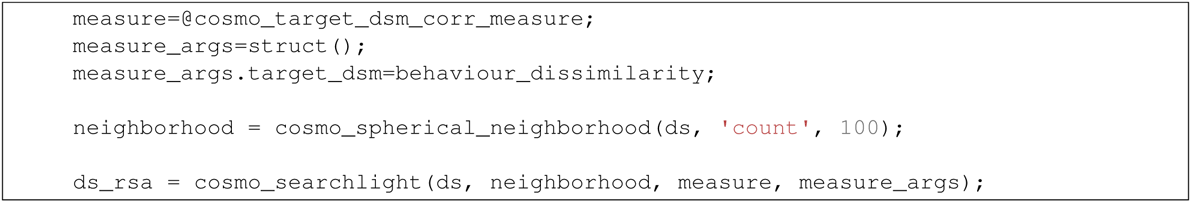

**Fig. 2:**
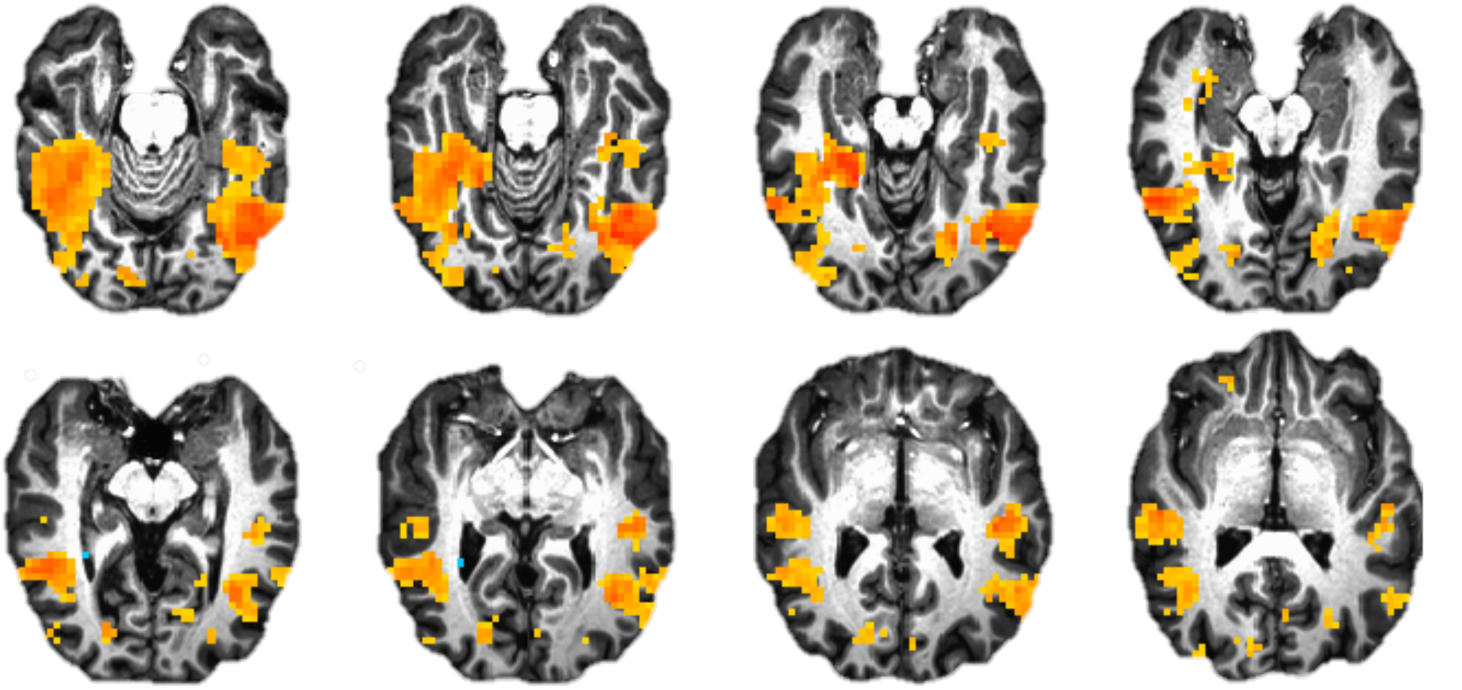
Illustration of representational similarity analysis searchlight results identifying regions where pattern similarity across six animal species is similar to behavioural similarity ratings.

For illustration purposes, the searchlight result map can be saved in a variety of volumetric formats:

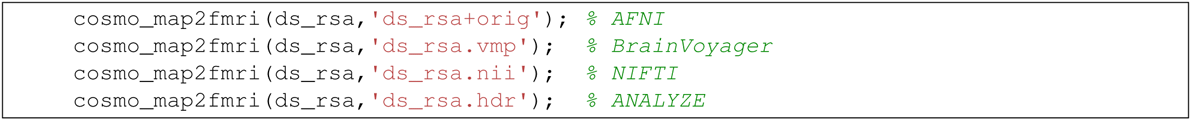

### 2.3 Locating effects in space and time

Using MEG data from a Neuromag306 system, gradiometer data is used to identify when and where 20Hz electrical median nerve stimulation can be identified using MEG data. Data from 145 trials were preprocessed in FieldTrip, selecting 145 epochs from trials during stimulation, and another 145 epochs from the same trials before stimulation. Each trial is considered to be independent from the others, and thus has a unique chunk value. As in the previous example, the only information that the user has to provide is the file to load, the targets (condition labels), and the chunks (indicating which samples are considered to be independent).

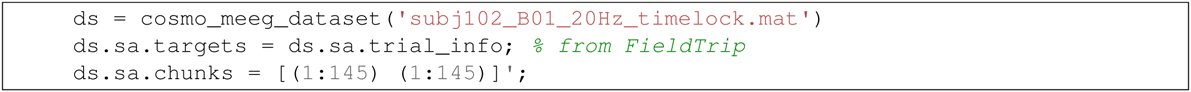

In the following analysis, a split-half correlation measure is applied (c.f. Haxby et al., 2001). Chunks are re-assigned so that about half of the samples have a value of 1 and the others a value of 2, allowing for computing correlations across the two halves.

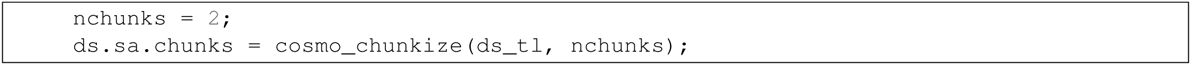

Similarly as illustrated in Figures 12 – 14, a channel-by-time neighborhood is defined by crossing a temporal with a spatial neighborhood. The input data is from pairs of planar gradiometers (two values per sensor location), whereas the output from the searchlight has combined gradiometer data (one value per sensor location).

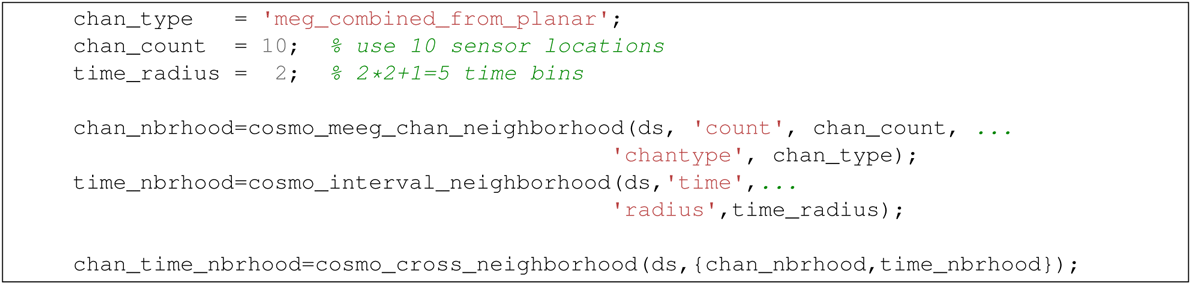

**Fig. 12:**
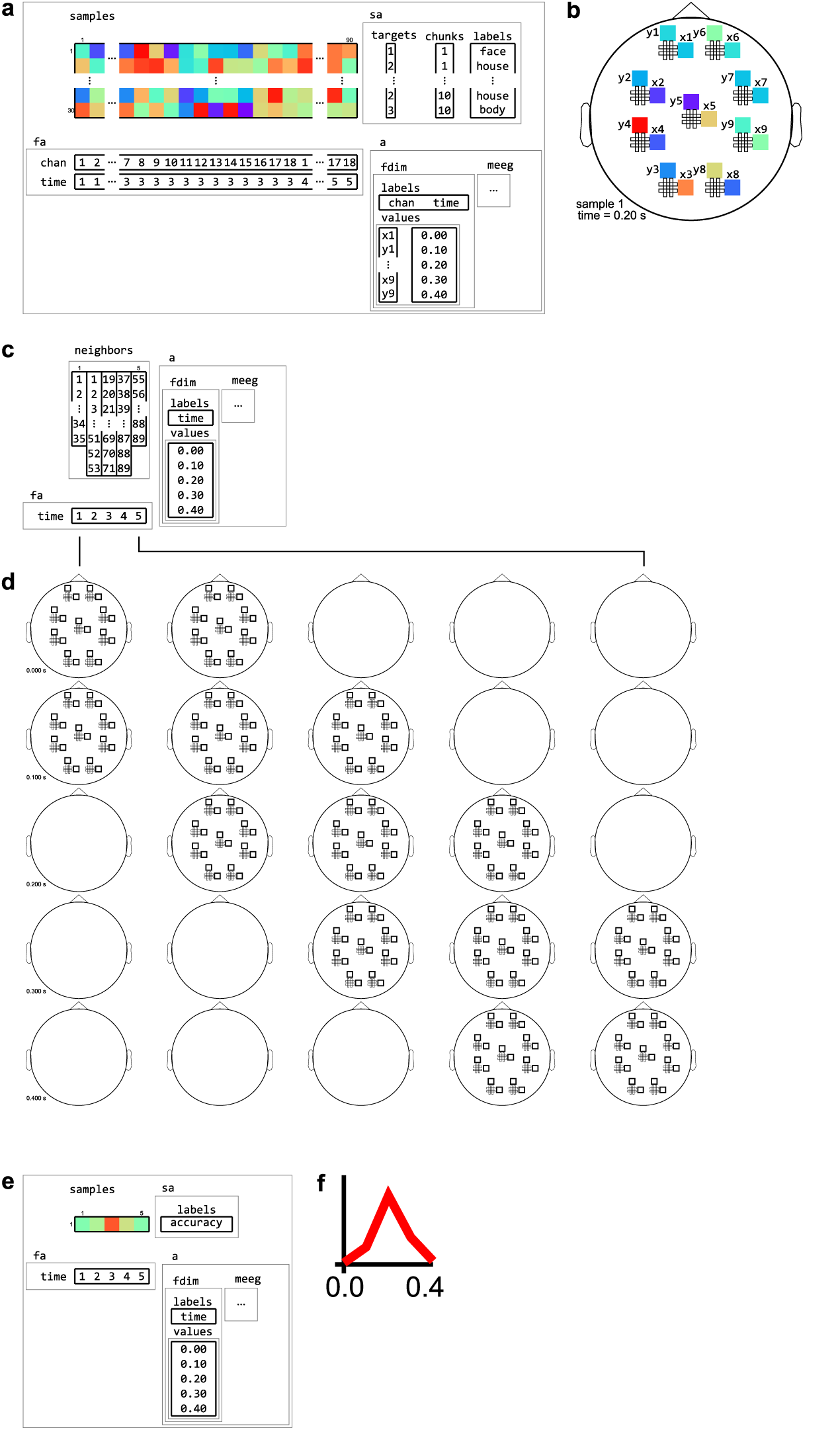
Example of neighborhood over time dimension. 1. A (tiny) M/EEG time-locked dataset with 3 conditions has, for each sample, data for 9 pairs of planar gradiometers (sensors `x1` to `x9` for one dimension, and `y1` to `y9` for the other one), and for 5 time points (from 0 to 0.4 seconds relative to stimulus onset). Its has two dataset dimensions, `chan` (sensors) and `time`. (b) Topology plot of data for the first sample (row) in the dataset, for the third time bin. (c) `cosmo_interval_neighborhood` produces a time interval neighborhood, with a radius of 1 time point. The output space of this neighborhood has a single dataset dimension `time`. (d) Illustration of neighboring features for each center feature in the neighborhood. For each center time point, its neighbors consists of features across all channels, and all time points that are at most one unit of time from the center. (e) Searchlight output when using the neighborhood defined in (c) with a cross-validation measure (not shown). The output has one sample (classification accuracy) and five features (one for each time point). (f) Schematic illustration of the dataset in (e), showing a time course of classification accuracies.

**Fig. 14:**
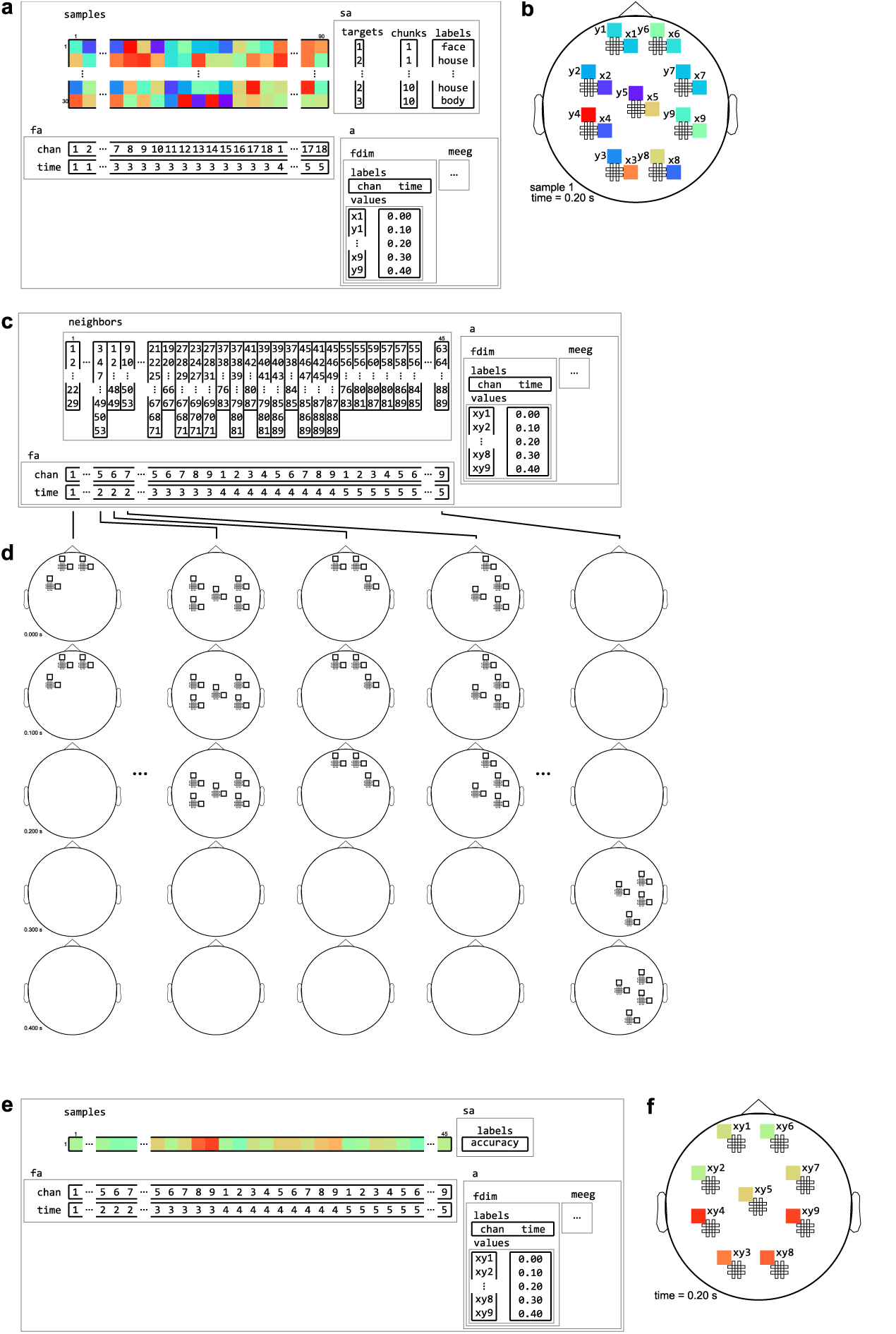
Example of neighborhood over channel by time dimension. 1. The same M/EEG dataset as in Figures 12 and 13, with two dataset dimensions, `chan` (sensors) and `time`. (b) Topology plot of data for the first sample (row) in the dataset, for the third time bin. (c) `cosmo_cross_neighborhood`, which crosses to the time neighborhood from Figure 12 and the channel neighborhood from Figure 13, produces a channel interval neighborhood, that selects neighboring channels within a certain spatial radius. The output space of this neighborhood has two dataset dimensions, `chan` and `time`. (d) Illustration of neighboring features for each center feature in the neighborhood. For each combined gradiometer and time point in the neighborhood, its neighbors consists of all time points that are at most one unit of time from the center, and for which the planar gradiometers are within a certain radius. (e) Searchlight output when using the neighborhood defined in (c) with a cross-validation measure (not shown). The output has one sample (classification accuracy) and 45 features (one for each pair of time points and combined gradiometers). (f) Schematic illustration of the dataset in (e) for a single time point. For each time point, there is a topology of classification accuracies for each combined planar gradiometer.

Using a searchlight and the `cosmo_correlation_measure`—with default options, i.e. the measurement argument is not required—each of the three neighborhoods can be used to produce a different searchlight map, locating effectors over time (across all channels), channels (across all time points), respectively. The last searchlight map, which contains a channel-by-time representation, is converted to a FieldTrip structure for visualization (see Figure 3).

**Fig. 3:**
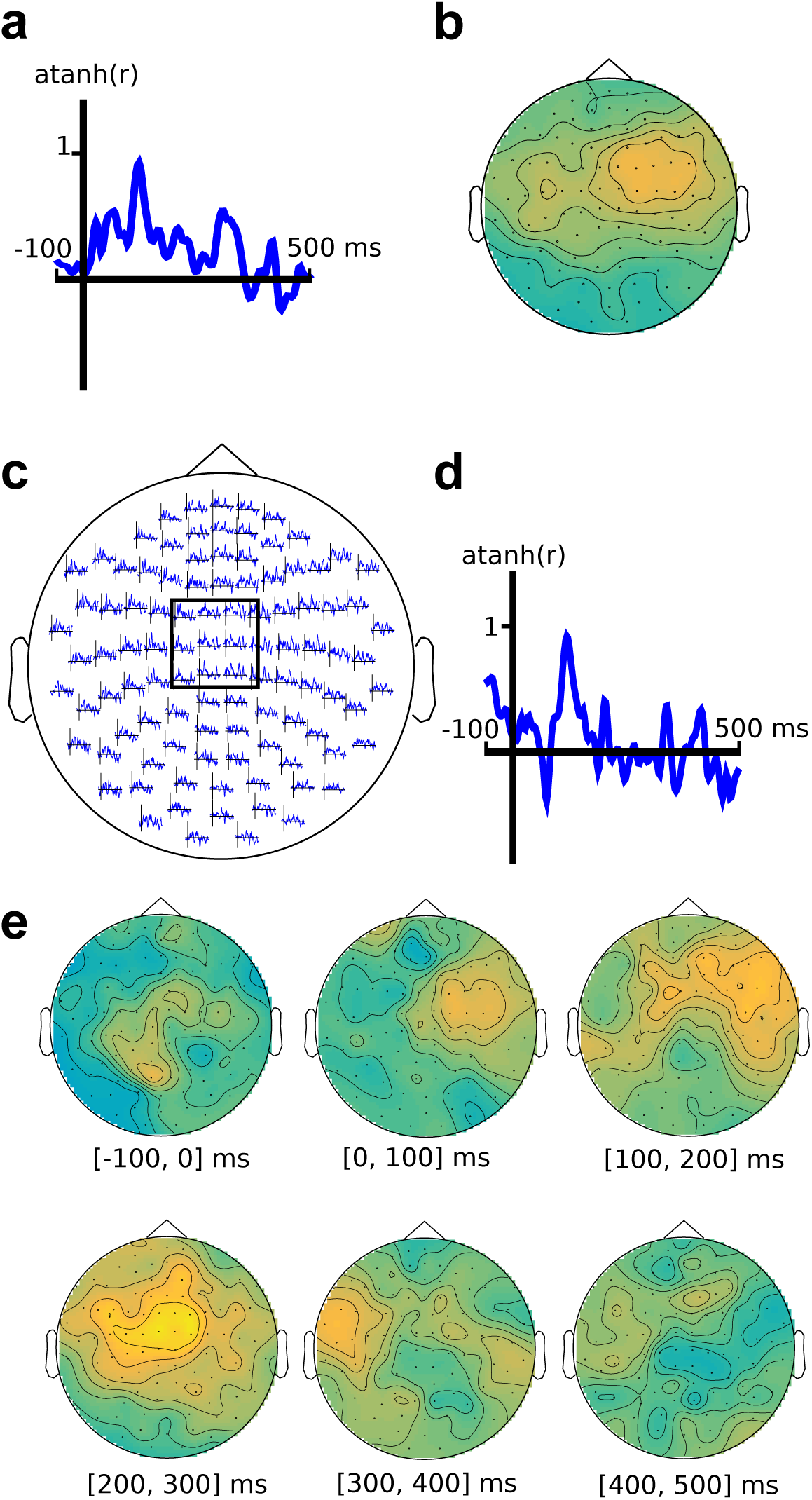
M/EEG searchlight maps in the time dimension, channel dimension, and channel-by-time dimensions. (a) time-only searchlight results using split-half Fisher-transformed correlation differences between somatosensory stimulation and rest. Each pattern contains data from all M/EEG channels. (b) channel-only searchlight using the same data as (a), where each pattern contains data from all time points. (c) channel-by-time searchlight, where each pattern contains data from a subset of channels and a subset of time points, with the inset (d) showing the average over a subset of sensors. (e) Alternative visualization of the results in (e), using topography plots for six time intervals.

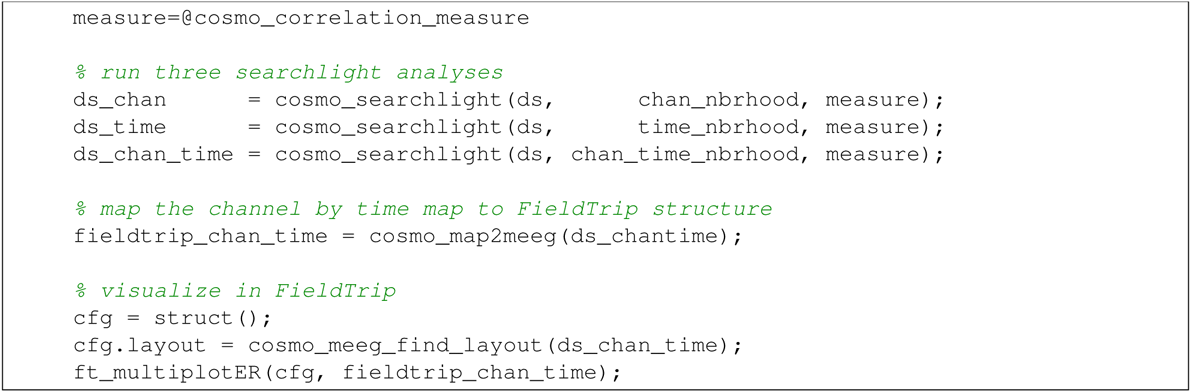

For illustration purposes, data is saved in FieldTrip and EEGLAB formats:

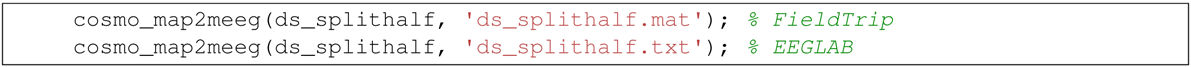

### 2.4 Generalization over time

It is also possible to measure pattern similarity across time. CoSMoMVPA provides `cosmo_dim_generalization_measure`, which implements the time generalization method (King & Dehaene, 2014). This function can use another measure (typically `cosmo_correlation_measure` or `cosmo_crossvalidation_measure`) to compute, for each pair of time points in a training and test set, correlations or classification accuracies. Train and test data are indicated by setting the `chunks` to 1 or 2, respectively. Because the function itself is a measure, it can be used with a searchlight, so that generalization over time can be located in space.

Using the same MEG dataset as in the previous example (which is channel-by-time along the feature dimension), first the time dimension is made a sample dimension. The resulting dataset has `time` along the sample dimension and `chan` along the feature dimension.

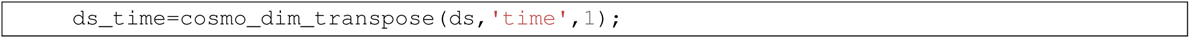

For the searchlight, a neighborhood along the chan dimension is defined as:

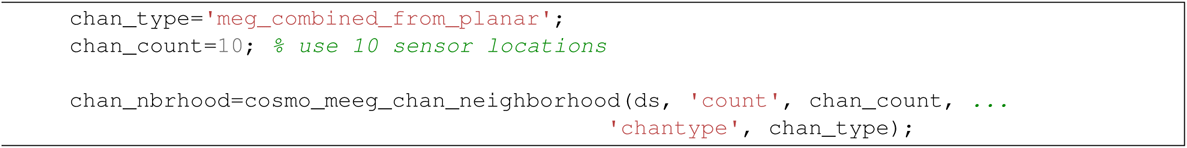

The measure uses an LDA classifier and another measure (`cosmo_crossvalidation_measure`) to compute classification accuracies. Time points are selected within a radius of `1` time points, so that for each pair of time points, generalization is computed across patterns spanning the current, preceding, and following time points.

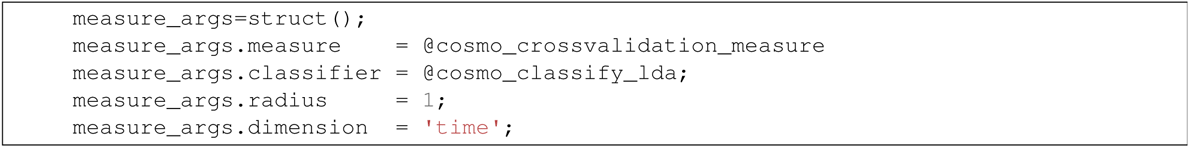

The searchlight is run similarly as in previous examples. The output has two sample dimensions, `train_time` and `test_time`.

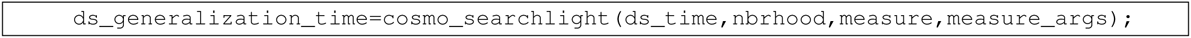

For visualization in FieldTrip, the sample dimensions are transposed into feature dimensions. As FieldTrip supports visualization of time-frequency data, the `train_time` and `test_time` dimensions are renamed to `freq` and `time`, so that FieldTrip is ‘tricked’ into treating the data as time-frequency data and can visualize the data for interactive exploration (see 4). FieldTrip can visualize the data directly:

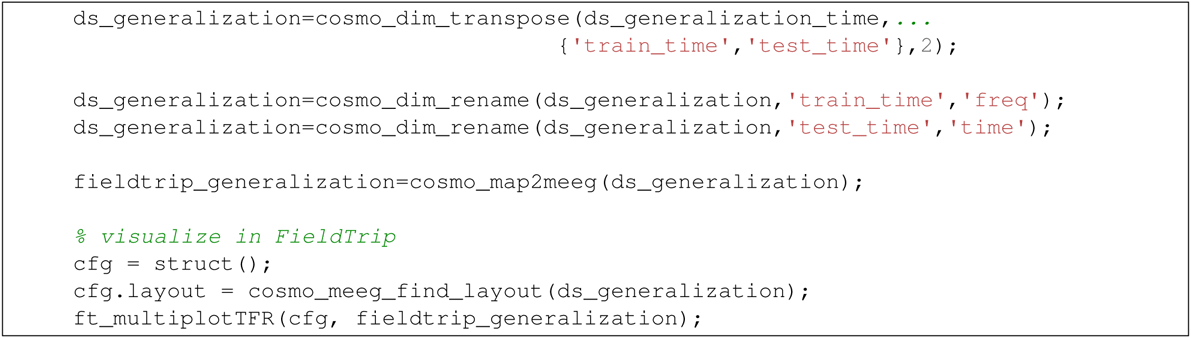

The next section describes the concepts used in CoSMoMVPA that underly the examples in this section.

## 3 CoSMoMVPA concepts

As we aimed to illustrate in the previous section, we believe that CoSMoMVPA provides an intuitive environment that is accessible to non-expert programmers using common data structures and interfaces used throughout the toolbox. The core concept is a common dataset structure (see Figure 5) inspired by PyMVPA (Hanke, Halchenko, Sederberg, Hanson, et al., 2009; Hanke, Halchenko, Sederberg, Olivetti, et al., 2009) extended with dimensionality information (Figure 6) to support various types of fMRI and M/EEG data uniformly.

**Fig. 5:**
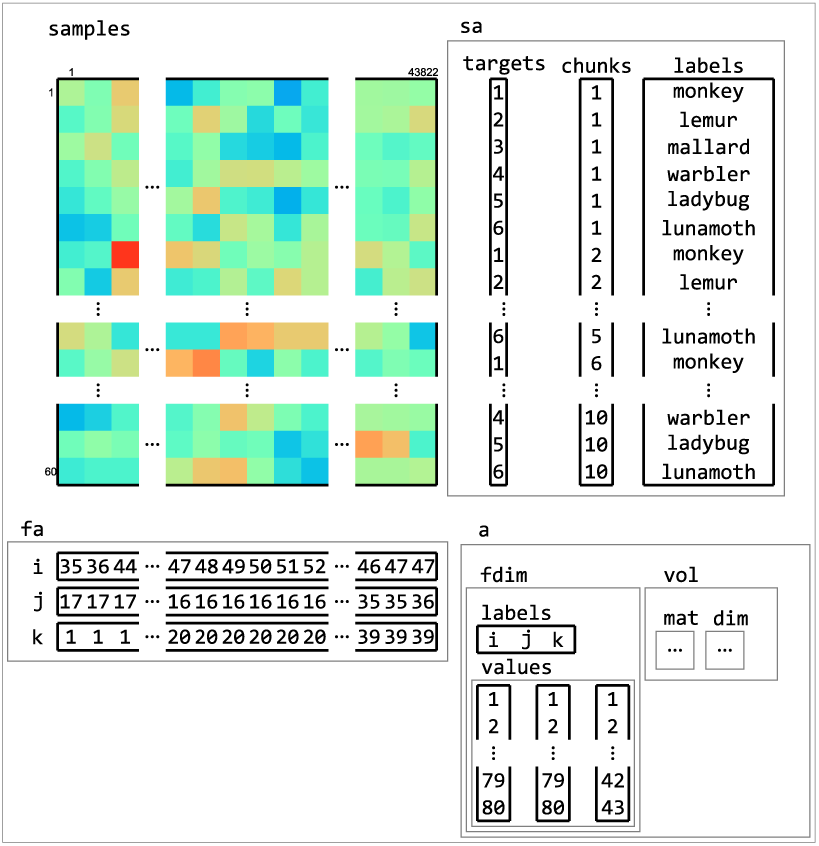
Example of dataset structure. This fMRI dataset has a samples matrix (top left) with 43882 features (voxels) and 60 samples (volumes). Sample attributes (`sa`, top right) contain information about conditions (in `targets`) and which samples are considered independent (in `chunks`, which, in this case, contains the acquisition run number). An additional sample attribute `labels` contain a human-readable description of each sample. Feature attributes (`fa`, bottom left) contain indices to the location of each feature. Dataset attributes (`a`, bottom right) contains information about the feature space of the dataset, which is used to determine neighbors of features (in searchlight analyses) and when exporting data to neuroimaging formats such as NIFTI.

**Fig. 6:**
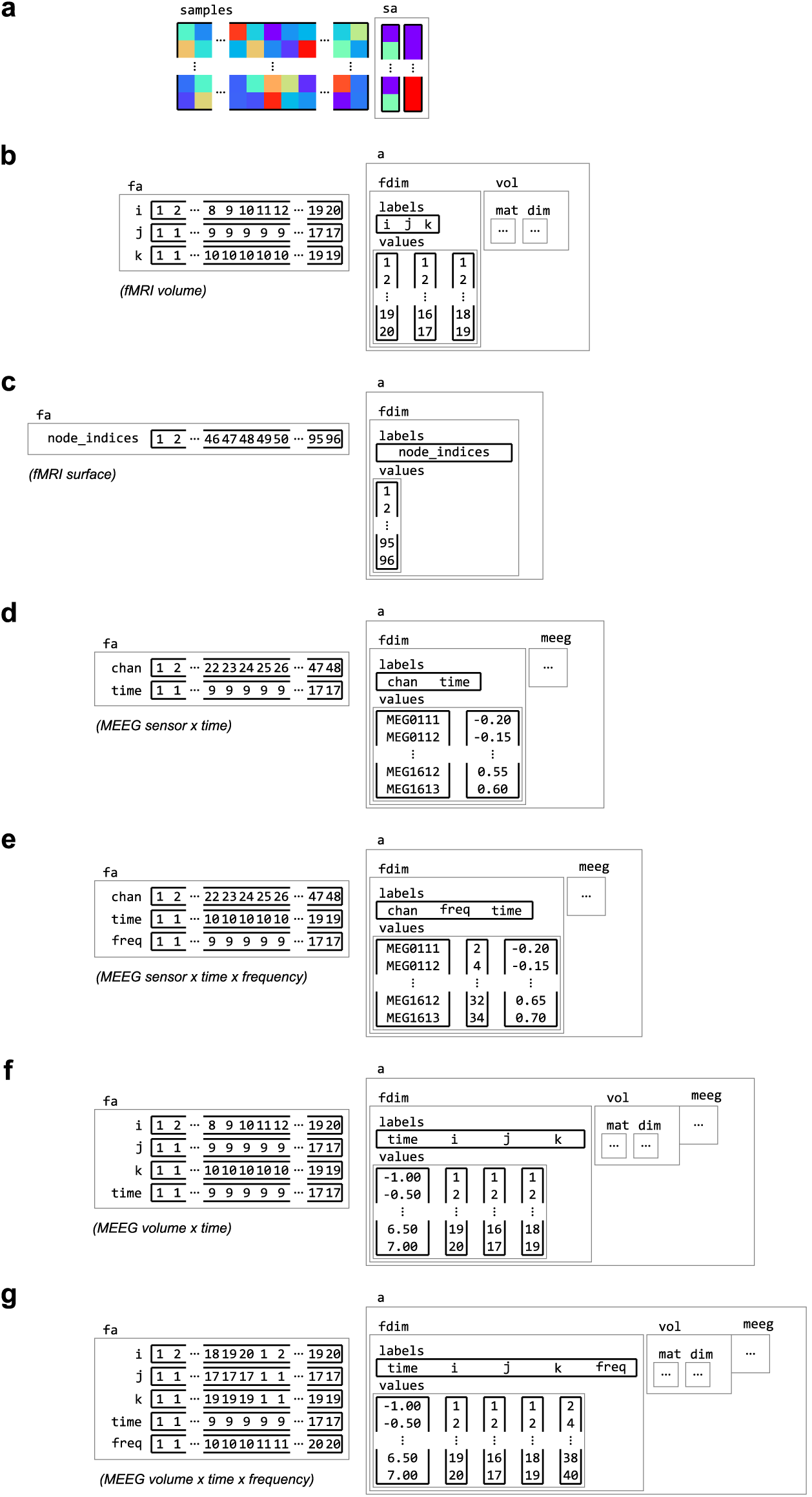
Examples of feature dimension attributes. Dataset samples and sample attributes (a) can be associated with feature and dataset attributes for various types of neuroimaging data with feature dimension information. Some examples of feature dimensions are for data in (b) fMRI volume, with three spatial dimensions `i, j` and `k`, (b) fMRI surface, with a single dimension `node_indices`, (c) M/EEG channel by time, with dimensions `chan` and `time`, (d) M/EEG frequency data, with dimension `chan, time` and `freq`, (e) M/EEG source data, which combines the spatial dimensions `i, j` and `k` with a `time` dimension, (f) and M/EEG source frequency data, with a frequency dimension `freq` added.

This common dataset structure allows for a variety of analysis types and dataset operations (Figure 8), including:

**Fig. 4:**
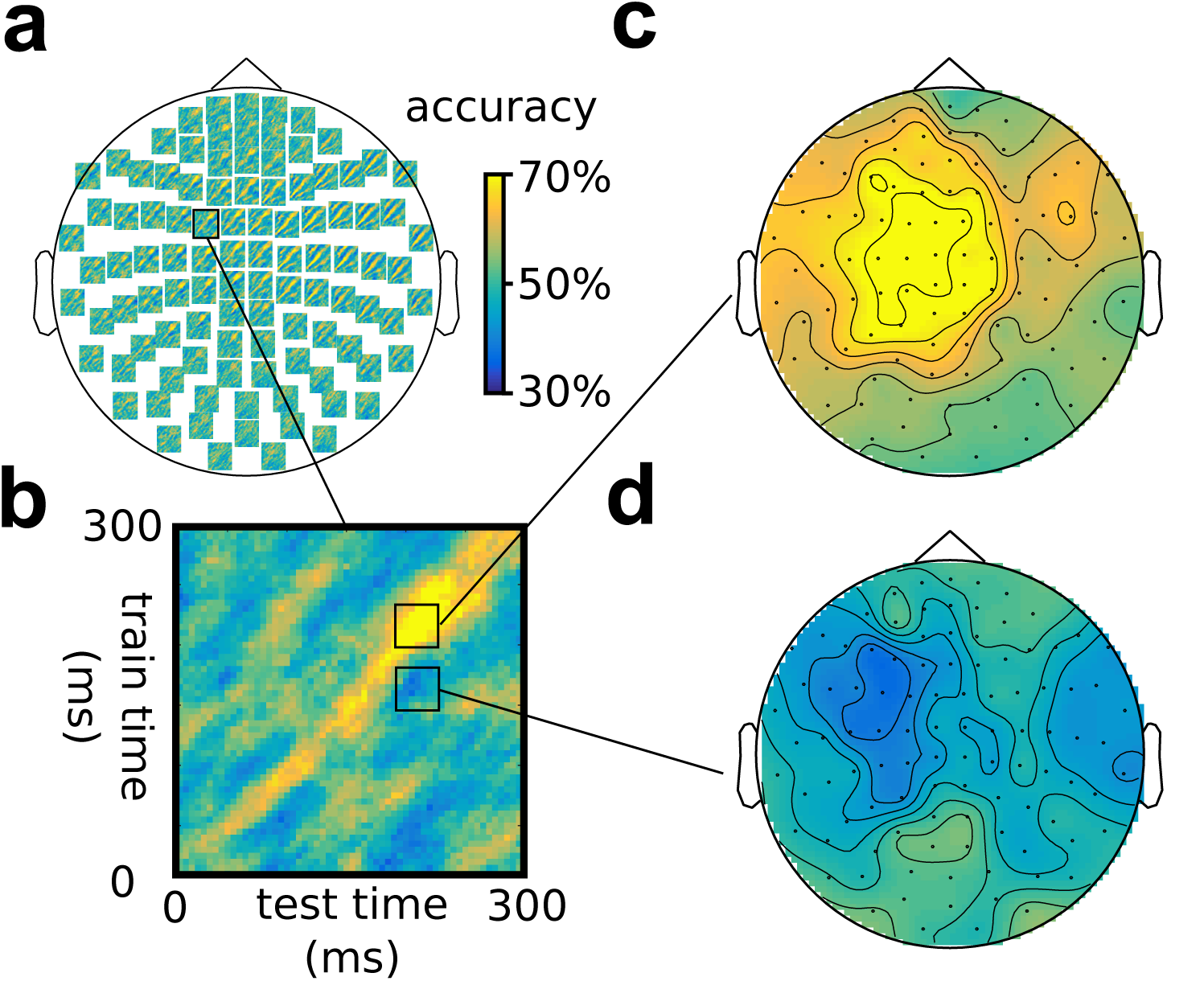
Illustration of generalization over the time dimension using an LDA classifier, using the same somatosensory stimulation data as in Figure 3. (a) Time-by-time accuracy plots for 102 combined gradiometers, with (b) inset of a single plot. (c, d) Topographies for two combinations of time intervals across the training and test time, showing above-chance accuracy on the diagonal (b) and below-chance accuracy off the diagonal (d).

**Fig. 8:**
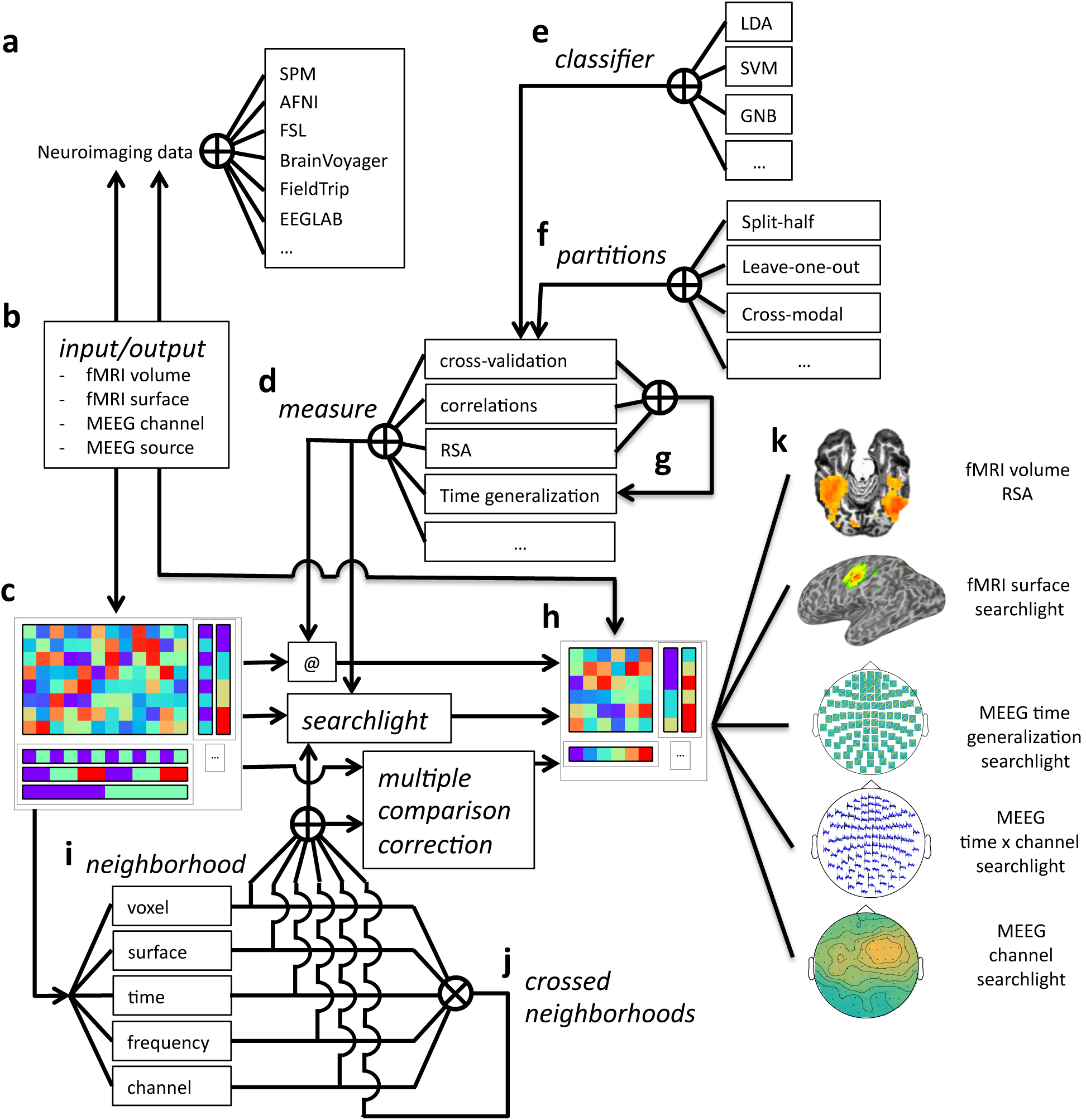
Overview of CoSMoMVPA architecture. Data collected (and optionally, preprocessed) in the fMRI or M/EEG modalities from a wide range of neuroimaging analysis packages (a) can be imported and exported (b) to a uniform CoSMoMVPA dataset structure (c). CoSMoMVPA provides several measures (d), including classification analysis (with various classifiers (e) and partitioning schemes (f)), correlation analysis, representational similarity analysis, and the time generalization method (which uses another measure for all possible pairs of time points for training and test sets (g)). Any measure can be applied directly (@) to the input dataset (c), which results in another dataset (h). CoSMoMVPA also provides various neighborhood definitions across voxel, surface nodes, time point, and frequency bin dimensions (h); these can be combined by crossing multiple dimensions (i). By combining a measure (d) with a neighborhood (i), a searchlight can be applied to the input dataset (c) resulting in another dataset representing a searchlight map (g). Neighborhoods are also used for Threshold-Free Cluster Enhancement multiple comparison correction. The result dataset from a measure, a searchlight analysis using a measure and neighborhood, or multiple comparison correction is a dataset (h) that, depending on the input and analysis parameters, can have various feature dimensions which can represent, for example, volumetric fMRI, surface-based fMRI, M/EEG topologies, M/EEG time series, M/EEG space by time data, or localized time generalization maps (j)-Result datasets can be converted back (b) for visualization or further analysis in a wide variety of neuroimaging analysis packages (a).

- merging and selecting subsets of datasets (Figure 9);
- conversion between matrix and multi-dimensional array representations (Figure 7) which is essential for input/output operations to fMRI and M/EEG file formats (Figure 10);
- defining MVPA measures with a common function signature, including split-half correlations (Haxby et al., 2001), classification predictions or accuracies using cross-validation with various partitioning schemes (D. D. Cox & Savoy, 2003), representational similarity analysis (Kriegeskorte, Mur, & Bandettini, 2008), and the time-generalization method (King & Dehaene, 2014);
- defining *neighborhood* structures in arbitrary feature spaces including time (Figure 12), space (Figure 13), and combinations such as time by space (Figure 14);
- *searchlight* analyses (Figure 15) using any type of MVPA measure and neighborhood structure.

**Fig. 7:**
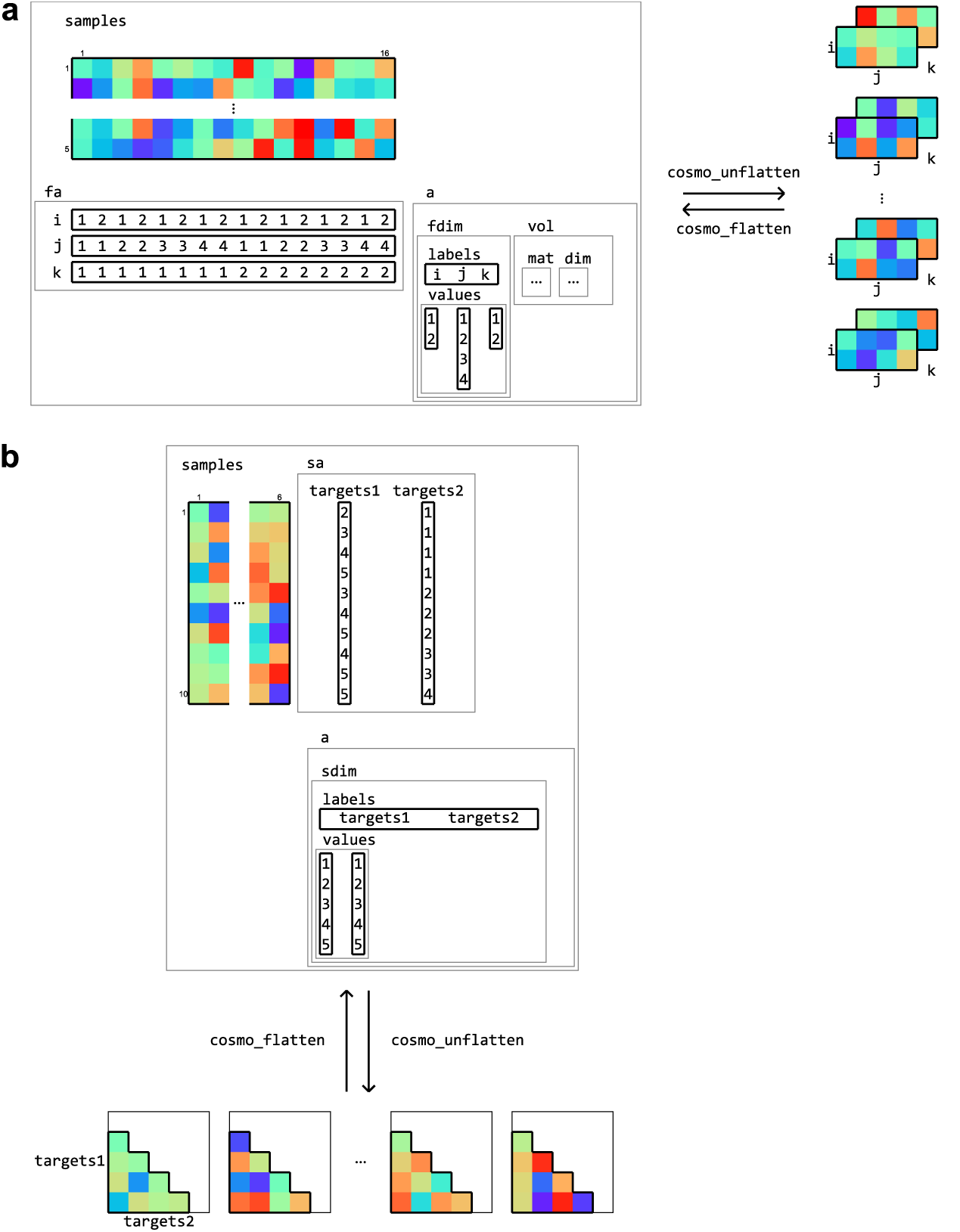
Flattening and unflattening datasets. A (tiny) fMRI dataset (a) with three spatial dimensions (left) is unflattened along the feature dimension using `cosmo_unflatten`. Each sample (row) in `samples` results in a three-dimensional array (right). (b) a dataset containing representational dissimilarities for five conditions is unflattened along the sample dimension. Each column in `samples` result in a dissimilarity matrix. The result from unflattening a dataset can be reversed in both (a) and (b) using `cosmo_flatten`. Note that dataset input/output functions for volumetric, surface-based, and M/EEG data (Figure 10) use the flattening and unflattening operations internally.

**Fig. 9:**
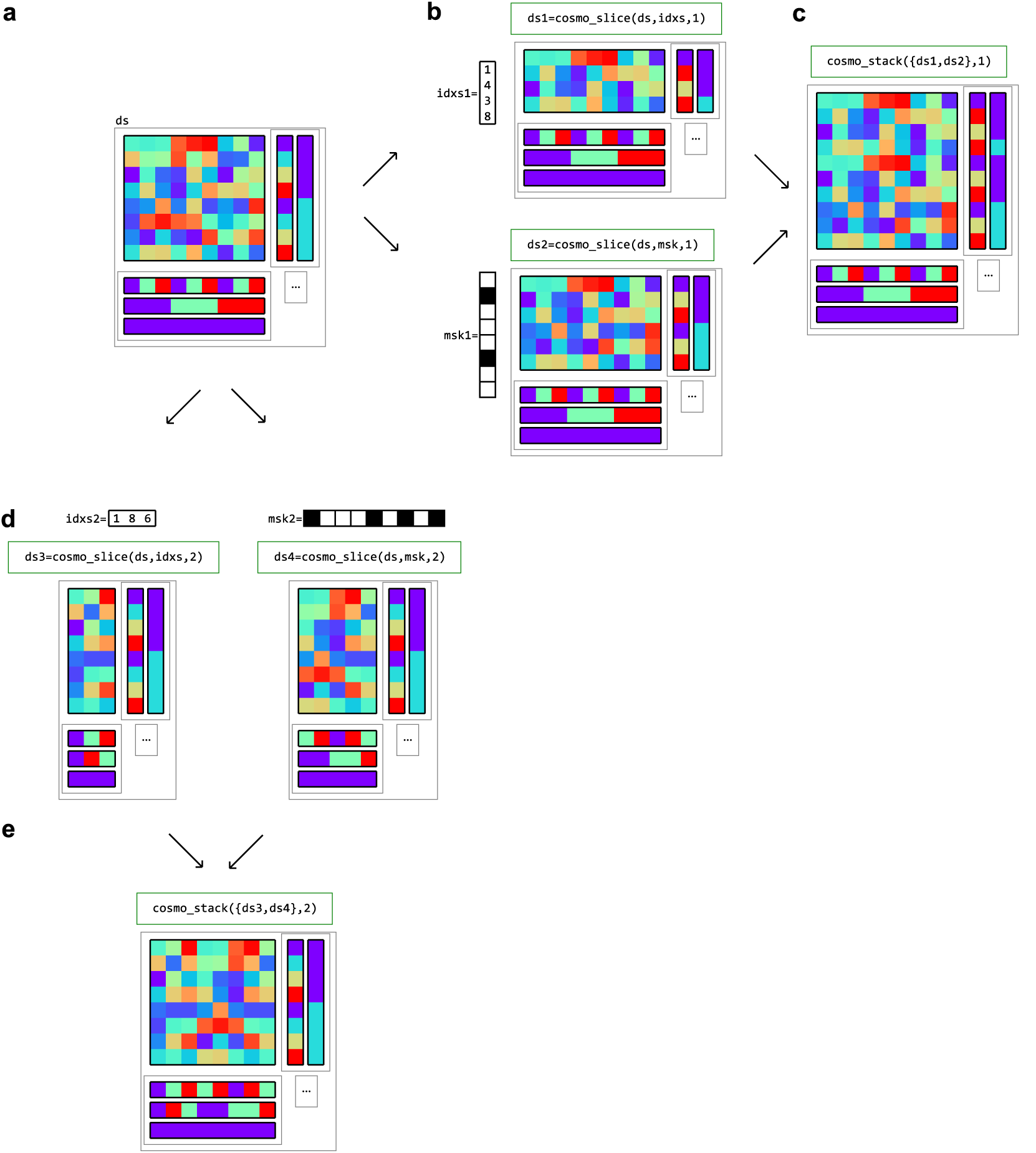
Example of dataset operations along the sample (first) dimension and feature (second) dimension. (a) a dataset structure can be sliced along the first (sample) dimension (b), using either indices (top) or a logical mask (bottom). The result has the same feature (`fa`) and dataset attributes (`a`) as the input, but only a subset of the samples and corresponding sample attributes (`sa`). One application is selecting data in a region of interest (ROI). (c) multiple datasets can be stacked along the sample dimension if the feature and dataset attributes match; one application is combining predictions after cross-validated classification analysis. (d) The same dataset can also be sliced along the second (feature) dimension. A typical use case is selecting a subset of samples for cross-validation analysis. (e) Multiple datasets can be stacked along the feature dimension if the sample and dataset attributes match; one application is combining results from different locations in searchlight analysis.

**Fig. 10:**
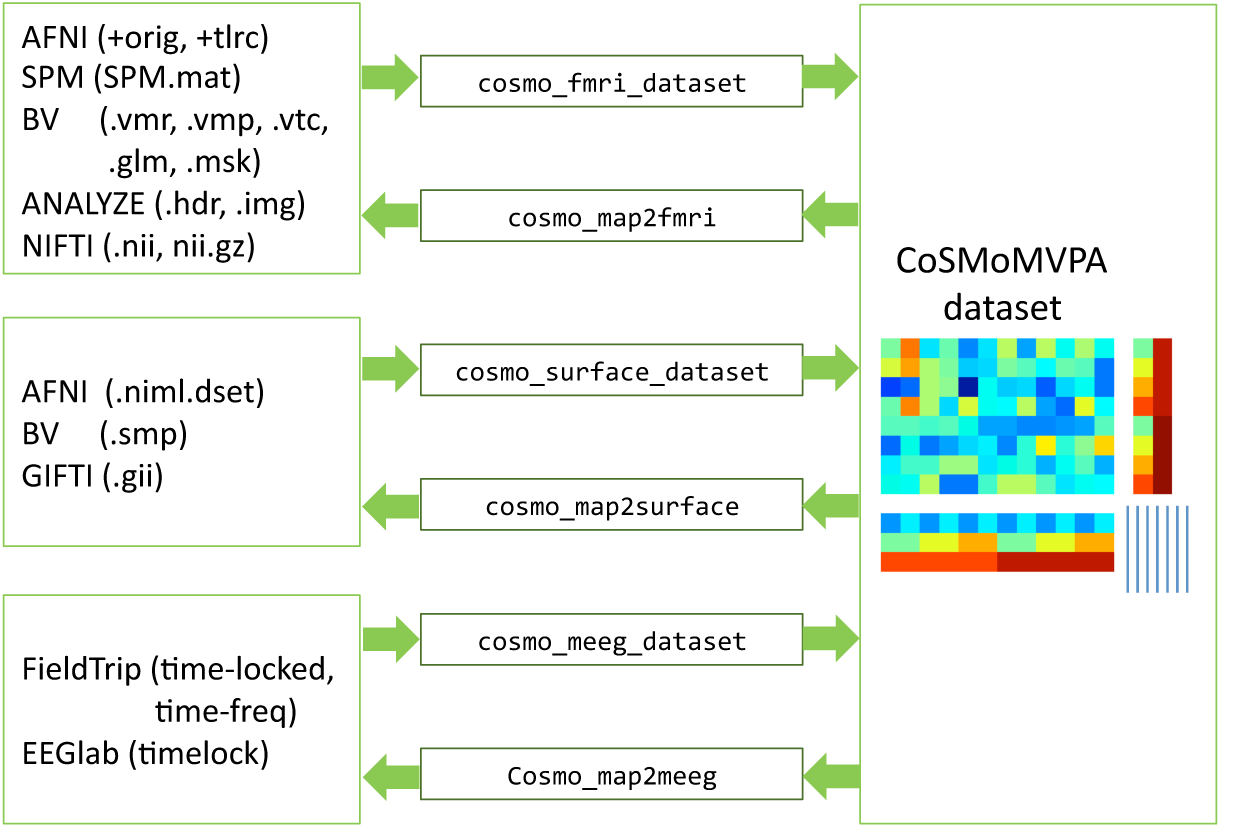
Dataset input/output support fMRI volumetric, fMRI surface-based, and M/EEG neuroimaging data (left) can be read and written by CoSMoMVPA through various input/output functions (center), and represented by a uniform dataset structure (right).

**Fig. 13:**
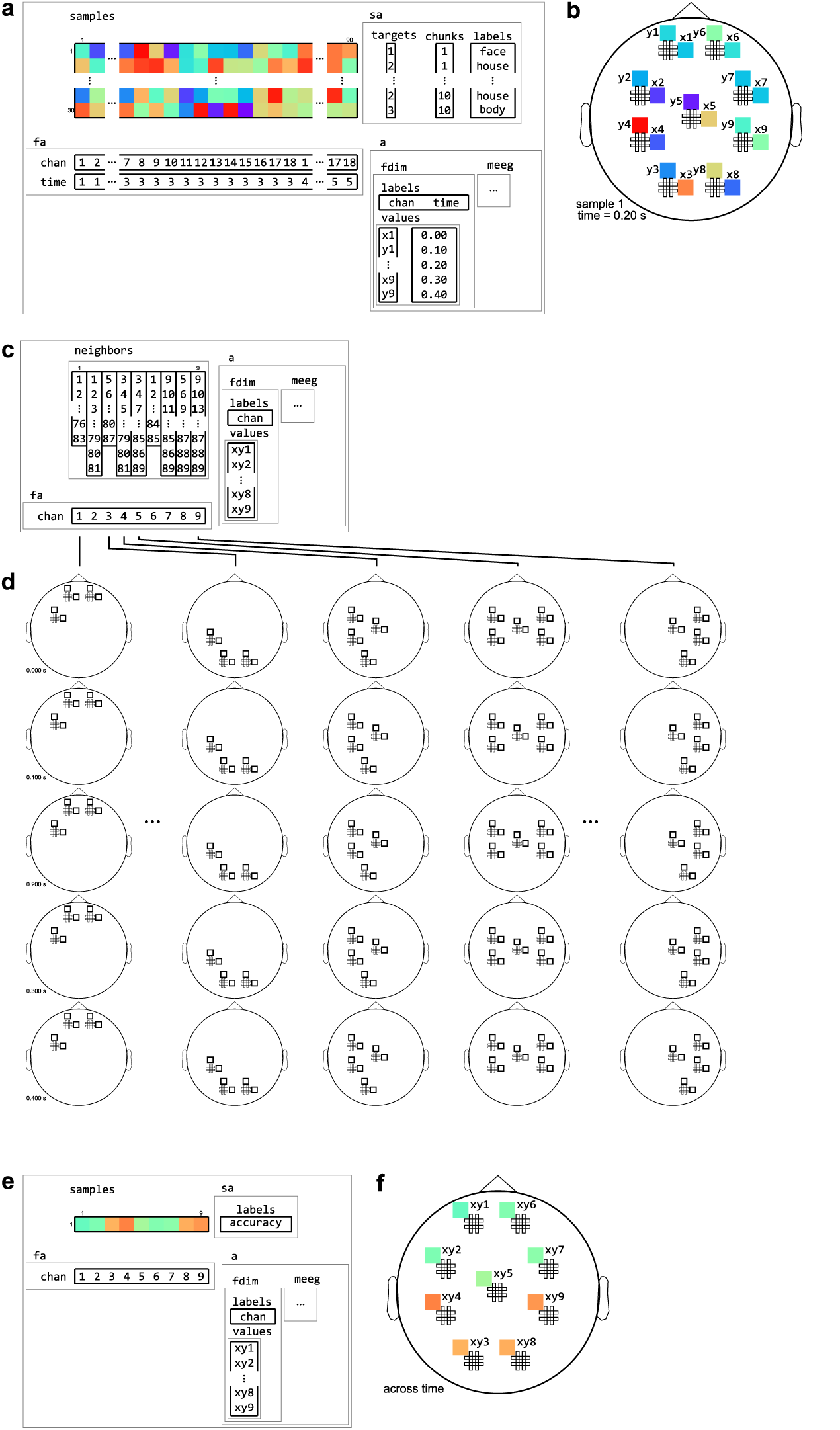
Example of neighborhood over channel dimension. 1. The same M/EEG dataset as in Figure 12, with two dataset dimensions, `chan` (sensors) and `time`. (b) Topology plot of data for the first sample (row) in the dataset, for the third time bin. (c) `cosmo_meeg_chan_neighborhood` produces a channel neighborhood that selects channels within a certain spatial radius. The output space of this neighborhood has a single dataset dimension `chan`. Note that the input dataset has 9 pairs of planar gradiometers making 18 sensors in total, whereas the neighborhood has 9 single ‘combined’ gradiometers. (d) Illustration of neighboring features for each center feature in the neighborhood. For each combined gradiometer in the neighborhood, its neighbors consists of all time points, and all planar gradiometers that are within a certain radius. (e) Searchlight output when using the neighborhood defined in (c) with a cross-validation measure (not shown). The output has one sample (classification accuracy) and nine features (one for each combined gradiometer). (f) Schematic illustration of the dataset in (e), showing a topology of classification accuracies for each combined planar gradiometer.

**Fig. 15:**
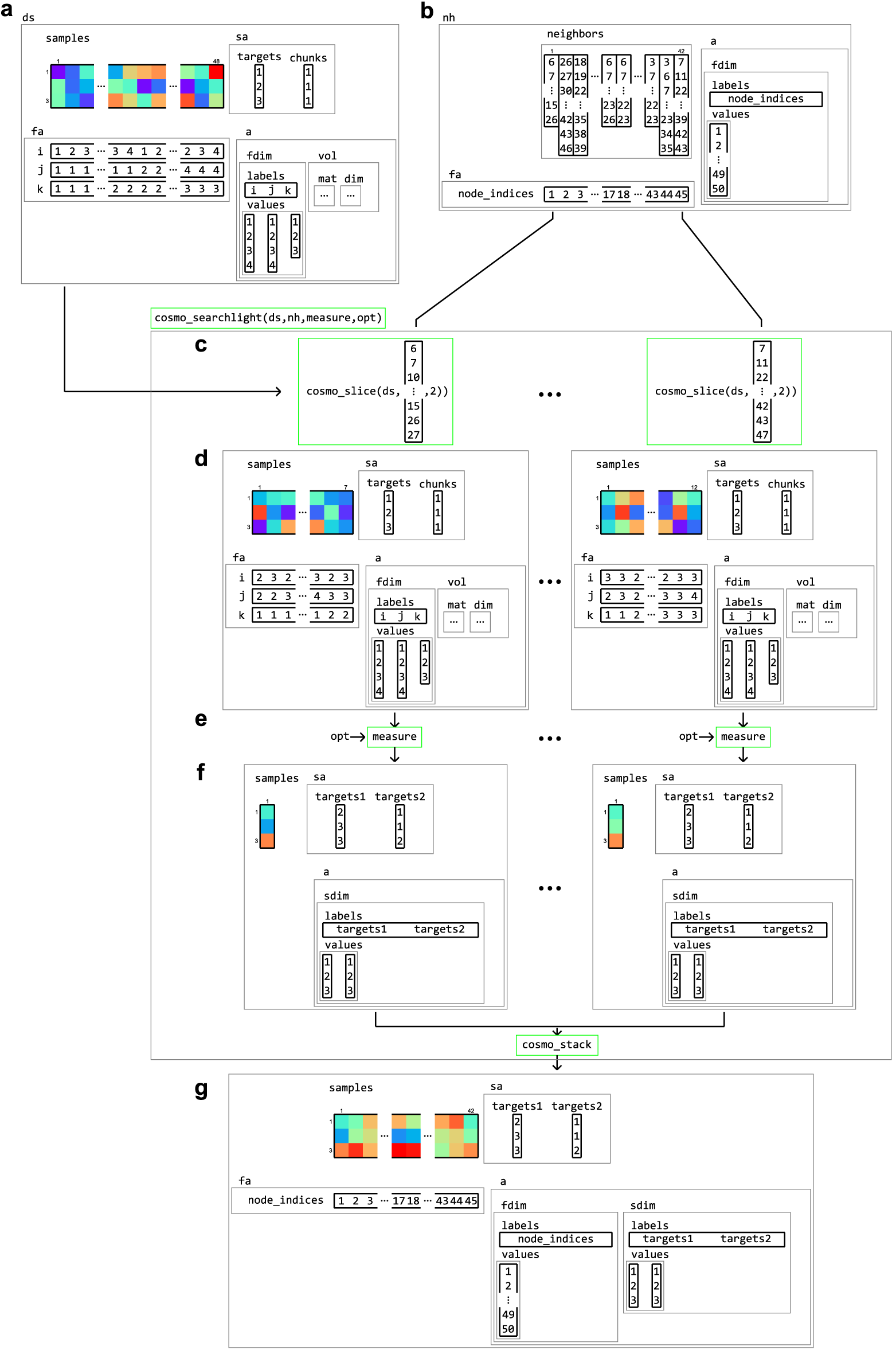
Example of volume-to-surface searchlight using a dataset structure, a neighborhood, and a measure. A (tiny) volumetric dataset (a) contains response patterns to three conditions (targets) and a single chunk. Using the dataset and surface models (not shown), a surface-based neighborhood (b) is constructed. Note that the feature attributes for the input volumetric dataset contain voxel indices, whereas the surface-based neighborhood contains node indices. A surface-based searchlight map is based on a dataset `ds`, a neighborhood `nh`, a function handle to a `measure`, and (optionally) options `opt` for the measure. Each searchlight center location is associated with an element in `neighbors`, containing a set of feature indices in the input dataset. Using these indices, the dataset is (c) sliced, resulting in a (smaller) sliced dataset (d) which is used as the input to a `measure`. Applying the measure results in an output dataset (f) associated with a single center location, which (here) contains pair-wise distances between samples in the input dataset. After repeating this for each center location, the output datasets (f) are stacked along the feature dimension and combined with the feature attributes `fa` and dataset attributes `a` from the neighborhood. The result (g) is a surface-based dataset that contains all pair-wise distances between patterns in local regions around each node on the surface from the input dataset.

In the remainder of this section, the CoSMoMVPA concepts underlying these analses types and operations are explained in detail.

### 3.1 Dataset structure

Inspired by PyMVPA, CoSMoMVPA uses a single dataset structure for all types of neuroimaging data (Figure 5). A minimal dataset has just one field `.samples`, which contains a samples-by-features matrix. Each row in the matrix corresponds to a *sample*, or observation; for example, these can be the blood-oxygen-level dependent (BOLD) response for a single volume, a *beta* estimate or *t*-statistic for a condition of interest during a single run, or the M/EEG signal during a particular time point during a single trial. In the context of CoSMoMVPA, each sample (row) is a pattern vector. Each column in the matrix corresponds to a feature, which refers to all measurements made for a particular location in an abstract ‘feature space’—where values at that location may be simple or derived measurements corresponding to some source. For example, features may correspond to voxels for volumetric data (M/EEG source and fMRI data), nodes (surface-based data), combinations of time points and M/EEG sensors (time-locked M/EEG data), or combinations of time points, frequency bands, and M/EEG sensors (M/EEG time-frequency data).

A dataset structure can contain additional attributes associated with the `.samples` field. Sample attributes (in a field `.sa`) contain information for each sample (row), and have the same number of rows as the `.samples` matrix. For most analyses, two sample attributes are required: `.sa.targets` and `.sa.chunks`. `.sa.targets` are used to store information about the experimental condition. `.sa.chunks` are used to index which subsets of samples can be considered as independent—such as fMRI acquisition runs. Typically their values are set by the user, although in certain cases they are set automatically, e.g. when loading SPM data. In addition, an arbitrary number of other attributes can be added as well, which may contain, for example, human-readable labels or statistical information.

Feature attributes (in a field `.fa`) is used to store indices specifying ‘locations’ of each feature, such as voxel indices, time point, or frequency band.

General dataset attributes (in a field `.a`) contain general information that is not specific to either features or samples. This can include information about the spatial or temporal dimensions of the dataset. For example, an fMRI volumetric dataset contains the affine transformation matrix that maps voxel coordinates to world coordinates in a field `.a.vol.mat`.

### 3.2 Dataset dimensions

In many cases, neuroimaging data is inherently multi-dimensional along the feature axis. For example, fMRI volumetric data has three spatial dimensions, M/EEG time-locked data has a spatial (channel) and a time dimension, and M/EEG time-frequency data has a spatial, time and frequency dimension.

Information about such dimensions is stored in a feature dimension attribute `.a.fdim`, which contains two subfields for labels (`.a.fdim.labels`) and values (`.a.fdim.values`). The values contain, for each dimension, a list of strings or numbers indicating the locations along that dimension. The labels contain the names of each dimension.

Each label must be a sub-field in the feature attribute field `.fa`, so that the vectors in the feature attributes index the elements in the values sub-field. Figure 6 gives examples of feature dimensions for fMRI and M/EEG datasets.

Along the same lines, data can also be multi-dimensional along the sample axis. A typical case is representing a dissimilarity matrix that contains distances between all pairs of samples in a dataset (see Figure 15).

Data can be flattened and unflattened along the feature and sample dimensions (Figure 7).

### 3.3 Dataset operations

As explained in the remainder of this Section, data in a CoSMoMVPA dataset structure can be processed in a variety of ways (Figure 8). `cosmo_slice` can be used to select particular samples (rows) or features (columns) with the corresponding sample or feature attributes.

One application of slicing samples is classification using cross-validation, where a subset of samples is sliced to form a training set, and another (disjoint) subset of samples is sliced to form a test set. In a similar manner, split-half correlations can be computed by slicing the dataset twice—once for each half.

The main application of slicing features is feature-selection, such as when only data in a pre-defined region-of-interest (ROI) is to be used for MVPA. Such a region can comprise a subset of voxels, nodes, or M/EEG channels; and/or a subset of time points or frequency bins. It is also used in searchlight analyses (described below), which involves the repeated application of slicing features.

Datasets can also be merged along the sample and feature axes using `cosmo_stack`. For the feature dimension this is useful when combining results from different locations, such as in searchlight analyses. For the sample dimension, rows can be stacked when combining predictions from different folds in a cross-validated classification analyses. Figure 9 gives examples of slicing and stacking along sample and feature dimensions.

### 3.4 Dataset input/output

CoSMoMVPA comes with a set of input/output functions that read and write fMRI volumetric, fMRI surface-based, and M/EEG data; these functions use the `cosmo_flatten` and `cosmo_unflatten` functions internally. Through various external toolboxes (Shen, 2010; Saad & Chen, 1999; Weber, 2010; Flandin, 2008), a wide variety of neu-roimaging formats is supported directly (See Figure 10).

### 3.5 Classifiers

A classifier function takes a *training* set of patterns `samples_train`, each with a target (class label) associated with them in a vector `targets_train`, and a *test* set of patterns `samples_test`. A classifier returns a predicted target label for each sample in `samples_test` in a vector `samples_test`.

CoSMoMVPA supports commonly used classifiers, including Linear Discriminant Analysis (LDA) and Support Vector Machines (SVM). Some classifiers support additional options, such as kernel selection in LIBSVM (Chang & Lin, 2011).

In CoSMoMVPA, all classifiers use the same signature as the LDA classifier illustrated here:

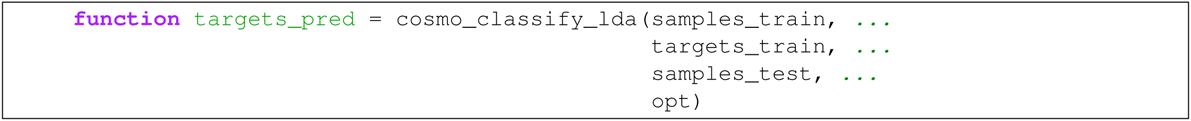

which, through the use of function handles, provides re-usable functionality for classification analysis when used with partitions and cross-validation measures. Note that this function signature makes explicit that the targets associated with the test set are not used for classification.

### 3.6 Partitions

Partitioning schemes, used for classifiation analysis, define which samples are used for training and for testing. They typically contain multiple folds to support cross-validation; in different folds, different samples are used for training and testing. CoSMoMVPA provides functions to generate a variety of partitioning schemes including those for cross-modal (Oosterhof, Wiggett, Diedrichsen, tipper, & Downing, 2010, Oosterhof, tipper, & Downing, 2012) and cross-participant (Clithero, Smith, Carter, & Huettel, 2011; Haxby et al., 2011) decoding (Figure 11). Using a cross-validation measure (explained below) with classifier provides functionality for cross-validation using a variety of cross-validation schemes and classifiers.

**Fig. 11:**
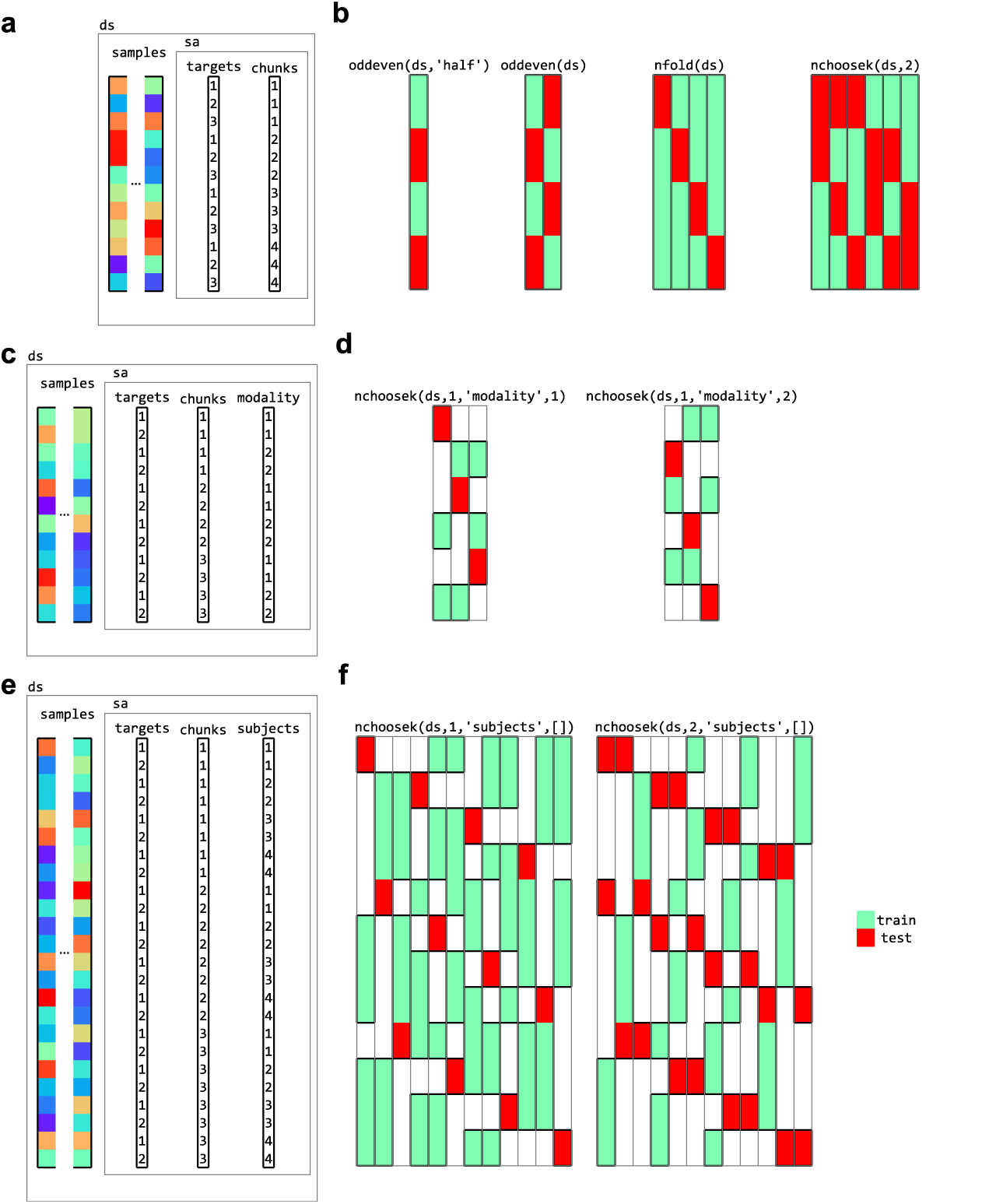
Partitions for cross-validation. (a) a dataset with three targets (conditions) and four chunks (for example, acquisition runs for fMRI data). Various cross-validation partitioning schemes can be defined using partitioning functions in CoSMoMVPA, including (b), from left to right, testing on even chunks after training on even chunks (using a single fold), testing on even chunks after training on even chunks and vice versa (two folds), testing on each chunk after training on the remaining chunks (four folds), and testing on each combination of two chunks after training on the remaining chunks (six folds). Each column represents a single fold, with green and red colors indicating samples used for training and testing, respectively. (c) a dataset with data acquired in two modalities (for example, visual and auditory). Cross-decoding cross-validation schemes can be defined for (d) leave-one-chunk out for testing, with testing on the first modality after training on the remaining chunks of the second modality (left) and vice versa (right). (e) a dataset with data from four participants, with (f) a cross-participant leave-one-chunk-out (left) and leave-two-chunks out (right) cross-validation schemes where data from one and two chunks, respectively, from one participant are used for testing after training on data from the other participants and the other chunks.

### 3.7 Dataset measures

In CoSMoMVPA, the *measure* concept is a generalization to compute one or more values of interest (such as classification accuracy, or pair-wise distances between patterns) from an input dataset. Measures have a common function signature: it takes an input dataset and optional arguments that prescribe the behaviour of the measure, and returns a dataset with a singleton feature dimension. Thus, its function signature is:

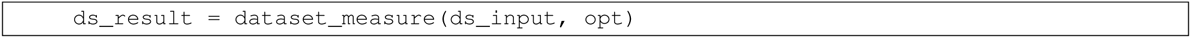

where `ds_result.samples` must be a column vector. CoSMoMVPA includes several measures for the most common MVP analyses, including correlations (Haxby et al., 2001), classification predictions or accuracies using cross-validation with various partitioning schemes (D. D. Cox & Savoy, 2003), representational similarity analysis (Kriegeskorte, Mur, & Bandettini, 2008), and the time-generalization method (King & Dehaene, 2014). In the case of region-of-interest analysis, measures can be applied directly to a dataset. As described below, when combined with a neighborhood, they can be used for generalized searchlights as well.

### 3.8 Neighborhoods

Together with measures, neighborhoods form a crucial ingredient for running searchlights (also known as information mapping), which provide a data-driven approach to localize multivariate effects of interest. The traditional searchlight (Kriegeskorte et al., 2006; Etzel, Zacks, & Braver, 2013) is used for volumetric fMRI data, where, for each ‘center’ feature (voxel) in the brain, other neighboring features within a certain radius, are used to compute a measure. The output from the measure is then assigned to that center feature.

More recently, surface-based searchlights (Oosterhof, Wiestler, Downing, & Diedrichsen, 2011; Chen et al., 2011) were proposed to improve feature selection—by selecting voxels in the grey matter—and use a more accurate anatomical delineation of cortical regions. In surface-based searchlights, each node is used as a center, and nearby voxels—in masks with shapes not dissimilar to curved cylinders—are used as input for MVPA; thus accounting for the folded nature of the cortical sheet. Because there is less “leakage” of information across sulci, informative regions can be localized more precisely and accurately (Oosterhof, Wiestler, Downing, & Diedrichsen, 2011). From an analysis perspective, in these searchlights one can distinguish between the input space, where the features are voxels in a volume, and the output space, where the features are nodes on a surface; whereas in the traditional searchlight, the input and output spaces are the same.

Through neighborhood structures, CoSMoMVPA generalizes the searchlight approach by supporting input and output spaces that may be different. A neighborhood structure contains, for each feature in the output space, a list of indices that map to features of the input space, as well as feature attributes and dataset attributes for the output space. Functions are provided to define neighborhoods in the volume (`cosmo_spherical_neighborhood`, for fMRI volumetric and M/EEG source space data), on the surface (`cosmo_surficial_neighborhood`), for M/EEG sensors (`cosmo_meeg_chan_neighborhood`), and over time and frequency bins (`cosmo_interval_neighborhood`). These neighborhood functions provide the following functionality:

- The volume and surface neighborhoods support selecting either features within a fixed radius, or, at each searchlight location, expanding the radius until a fixed number of features is selected.
- The surface neighborhood can be defined with input data either in volume or surface space.
- Interval neighborhoods can be defined by setting a one-dimensional ‘radius’ that defines how wide the bins are along the time or frequency dimensions. (A radius of zero means to only use each center time point or frequency bin itself as neighbor). An example is given in Figure 12.
- M/EEG sensor neighborhoods can be based on any channel layout supported by FieldTrip Oostenveld et al., 2011. This means that all commonly used EEG and M/EEG systems supported in FieldTrip (currently 32) are supported in CoSMoMVPA;.
- Certain systems, such as the Neuromag 306 system, use two types of sensors: gradiometers and planar gradiometers. Neighborhoods can be defined for either sensor type. In the case where planar gradiometers come in pairs, the input space consists of pairs of planar gradiometers, and the output space of single ‘combined’ gradiometers. The advantage is that the output maps have one value per sensor location, but that the input is based on the original individual gradiometer data without the need for averaging. An example is given in Figure 13.

Different neighborhoods can be crossed to form new neighborhoods. For example, an M/EEG channel neighborhood can be crossed with a time interval neighborhood to produce a channel-by-time neighborhood, which allows for locating effects of interest over space and time (see Figure 12). Similarly a spherical neighborhood in M/EEG source space can be crossed with a time interval neighborhood. When M/EEG data is in time-frequency space, a frequency interval neighborhood can be included as well, so that effects can be located over space, time, and frequency bin (c.f. Leske et al., 2014, Tucciarelli, Turella, Oosterhof, Weisz, & Lingnau, 2015). For fMRI data, the lower temporal sampling rate and low-frequency characteristics makes temporal inferences more challenging, but if a sufficient number of volumes are acquired per trial, a volume or surface neighborhood can be crossed with a time interval neighborhood to run a searchlight over space and time (for an example, see Linden, Oosterhof, Klein, & Downing, 2012, Supplementary Animation 1).

### 3.9 Searchlight

Based on the building blocks introduced earlier, a searchlight analysis becomes trivial. It requires a dataset structure, a neighborhood structure, and a measure. Running a searchlight analysis involves slicing the dataset according to the feature indices in the neighborhood structure, applying the measure to each sliced dataset, stacking the results, and adding feature and dataset attributes from the neighborhood structure to form the output. Since it is based on the general concepts of a neighborhood and measure concept, it can be used for any measure and any type of neighborhood. As illustration, the internal workings of an fMRI surface-based searchlight that computes the pair-wise dissimilarity matrix from neighborhoods consisting of voxels is illustrated in Figure 15.

### 3.10 Multiple comparison correction

For multivariate searchlight as well as univariate analyses, locating effects of interest based on statistical whole-brain feature maps (in the volume, on the surface, or in M/EEG sensor, time, sensor by time, voxel by time, sensor by time by frequency, or voxel by time by frequency) must take into account chance capitalization. That is, correction for false positives must consider the large number of statistical tests that are performed. Since the meaningful units in neuroimaging analysis are *clusters* of features (Penny, W & Friston, 2003), rather than individual features, the multiple comparison correction approach taken in CoSMoMVPA is cluster-based as well.

Certain approaches are less suitable for multiple comparison correction. First, Bonferroni correction remains conservative even under the assumption of independence, but as typical statistical maps from neuroimaging analyses are smooth, this results in correction that is far too conservative. Second, False Discovery Rate, is not suitable for controlling for false positives in clusters of features in neuroimaging data Chumbley & Friston, 2009.

Third, fixed-threshold cluster-based correction is also a popular method and available in all major neuroimaging packages. It requires defining an uncorrected feature-wise threshold first. Using some estimate of the smoothness of the data (e.g., AFNI, SPM, BrainVoyager) or through a sample randomization scheme (e.g., FieldTrip), a critical cluster size is computed, using either Monte Carlo simulations (e.g. AFNI, BrainVoyager, FieldTrip) or Random Field Theory (SPM). However, this method has one main disadvantage: the use of a fixed feature-wise threshold. Different thresholds may lead to entirely different surviving clusters. For example, if a map has one large cluster with moderately significant statistics, and another small cluster with very significant statistics, then using a low-significance threshold makes the first cluster survive but not the second, whereas a high-significance threshold has the opposite effect. Apart from increasing the researcher’s degrees of freedom, it can also lead to instability in clustering results, because small differences in the input can determine whether two clusters below the critical size are connected beyond the feature-wise threshold (and may survive correction) or not.

To address this, Threshold-Free Cluster Enhancement (TFCE; Smith & Nichols, 2009) computes an aggregate score for each feature after the statistical input map has been thresholded over a wide range of levels. It avoids the issues associated with a fixed-threshold approach, and also supports different smoothing levels of the data, allows for interpreting local maxima across a cluster with large spatial extent, and has been validated for both fMRI and M/EEG data (Mensen & Khatami, 2013; Salimi-Khorshidi, Smith, & Nichols, 2011).

CoSMoMVPA supports Threshold-Free Cluster Enhancement in combination with a neighborhood structure that is computed automatically based on input dataset dimensions. This means that multiple comparison correction can be achieved for all datasets with dataset dimensions supported in CoSMoMVPA, including data with volume, surface, source space, sensor space, time, and frequency dimensions; and combinations of those. Using such a neighborhood, Threshold-Free Cluster Enhancement through Monte Carlo simulations is supported at the group level through a function `cosmo_montecarlo_cluster_stat`. Appropriate randomized datasets can either be computed using sample-based randomization (as also implemented in FieldTrip; Maris & Oostenveld, 2007), or through randomization at the individual participant level (Stelzer, Chen, & Turner, 2012). For comparison purposes, there is also support for fixed-threshold cluster-based correction, although this method is not recommended.

Altogether, the building blocks presented here can be used for a wide variety of MVPA approaches for fMRI and M/EEG data, supporting commonly used measures and a generalized searchlight.

## 4 Design descisions

### 4.1 The Matlab / GNU Octave language

CoSMoMVPA is implemented in the intersection of the Matlab and GNU Octave languages. Using the Matlab / Octave platform means it can use several existing toolboxes to support a variety of input and output formats, and use surface-based neighborhood definitions through the surfing toolbox (Oosterhof, Wiestler, & Diedrichsen, 2011). Importantly, M/EEG data preprocessed in EEGLAB or FieldTrip can be imported in CoSMoMVPA, and MVPA results can be visualized in FieldTrip directly.

Matlab (and, to a lesser extent, GNU Octave), is a popular platform in cognitive neuroscience research, with many other widely used packages running on it. Examples include Psychtoolbox (Brainard, 1997) for running experimental paradigms, FieldTrip (Oostenveld et al., 2011) and EEGLAB (Delorme & Makeig, 2004) for M/EEG analysis and visualization, and SPM (Friston et al., 1994) for fMRI analysis and visualization. It is also widely used for general scientific data analysis. As a result, many researchers may already be familiar with the Matlab / GNU Octave language, which would limit the time investment to learn CoSMoMVPA.

Although many institutions provide Matlab licenses to their students and staff, a disadvantage of a Matlab-only toolbox (without GNU Octave support) is that (1) access to the license is required to use Matlab, and (2) the full internal workings of data processing cannot be studied in full detail. CoSMoMVPA does not have these disadvantages, which makes it available for those who do not have access to a Matlab license. Indeed, CoSMoMVPA runs on fully open source platforms such as the NeuroDebian distribution. From a scientific perspective, this means that the analyses performed by CoSMoMVPA can be studied, verified and reproduced in arbitrary detail. From a practical perspective, this makes it feasible to run analyses on a computing cluster with no limits on the number of processing cores used, or on cloud-based computing services such as Amazon web services. Another advantage, explained in more detail below, is that the software can be tested automatically using online services that run on open source software only.

At the same time, GNU Octave does (as of 2015) not support all features present in Matlab, and many toolboxes specifically aimed at the Matlab platform are not fully compatible with GNU Octave. This may limit adoption of this platform, although GNU Octave is under active development and it is expected that it will support, in the future, more features that are currently Matlab-specific.

It should be noted that Python is another popular programming language, which (unlike Matlab) is fully open source. PyMVPA provides some advanced functionality currently not present in CoSMoMVPA. To support advanced users who want to use such functionality, PyMVPA provides data interchange with CoSMoMVPA through a `cosmo` module. In particular, datasets and neighborhoods from CoSMoMVPA can be used directly in PyMVPA, so that all MVP functionality present in PyMVPA can be applied to CoSMoMVPA data (including searchlight analyses); results can be exported from PyMVPA and loaded in CoSMoMVPA. A particular use case is MVPA of M/EEG data, for which input/output support and neighborhood definitions in PyMVPA is limited.

### 4.2 Maintainable architecture

This section discusses the architecture of CoSMoMVPA, and is mostly aimed for those interested in the design and development of CoSMoMVPA. It can be skipped by those whose are only interested in using the software.

The design of the architecture forms the basis of software. It has been proposed (Spinellis & Gousios, 2009) that it is desirable for an architecture to follow the following principles / properties:

- versatility: offer “good enough” mechanisms to address a variety of problems with an economy of expression. CoSMoMVPA supports all commonly used MVPA analyses in the fMRI and M/EEG data domains, and typical analyses require only a few lines of code. Support for a wide variety of neuroimaging formats means that it can interchange data with a variety of packages for analysis and visualization. CoSMoMVPA uses a modular approach, following the Unix philosophy that each function should do one thing, and do it well. Functions serve as building blocks that can be combined for complex analysis pipelines.
- conceptual integrity: offer a single, optimal, non-redundant way for expressing the solution of a set of similar problems. Through a common dataset and neighborhood structure across fMRI and M/EEG datasets, and uniform classification and measure function signatures. This solves the general problem of how to represent data across a variety of dimensions in space (fMRI and M/EEG voxels, fMRI node indices, and M/EEG channels), time, and frequency bins.
- independently changeable: keep its elements isolated so as to minimize the number of changes required to accommodate changes. Functions in CoSMoMVPA that implement particular classifiers, measures, neighborhoods can be changed independently, if necessary. For example, if one were to change how data is normalized when used in a representational similarity analysis searchlight, then only changes in the measure are required, not in the searchlight code itself, or the neighborhood functionality. The user can also implement their own classifiers, measures, or neighborhoods, which can be incorporated directly with the existing functions to support region-of-interest or searchlight analyses.
- automatic propagation: maintain consistency and correctness, by propagating changes in data or behavior across modules. The source code for documentation, code examples, and exercises is maintained in a single place. The documentation build system uses Sphinx-doc (Brandl et al., 2008) and Sphinx extensions (Mikofski, 2009, Troffaes, 2011), so that documentation and the output from running code examples is translated into HTML format automatically—its output is used as the content of the website. The build system supports exercise code, which is standard Matlab / GNU Octave code with special comment tags % >@@> and % <@@<< around lines to be used as an exercise. Such code is transformed into two versions: a full version with all the code, and a skeleton version which has the code in between the tags replaced by a comment stating ‘your code comes here’. This facilitates writing exercises and ensures that the full exercise code is consistent with skeleton code. The use of the build system ensures that the contents of the HTML documentation and website is consistent across documentation, code examples, and exercises.
- growth accommodation: cater for likely growth. Using the `git` version control system and `github` to coordinate distributed development means that contributions can be reviewed and, if quality standards are met, included in the code base through a convenient web interface. Through `git`, any modification in the code is recorded, and for any file its changes over time, and who made the changes, are tracked and can be reported. The use of `git` and `github` for distributed development has been, and is currently, used successfully in the PyMVPA project. Most software packages are not static: over time, more functionality is added, and bugs are fixed. CoSMoMVPA uses a modular architecture that we believe is easily extensible to include support for new functionality, including data formats, classifiers, and measures.
- entropy resistance: maintain order by accommodating, constraining, and isolating the effects of changes. Through the lifetime of a project, as functionality is added, improved or changed, entropy will increase, for example when the same functionality is implemented, over time, in various modules, leading to code duplication. Code duplication leads to more maintenance efforts, and can lead to inconsistencies if changes are applied in one place but not others. Entropy can also increase when new functionality is added incrementally for specific use cases, even if such use cases can be expressed more succinctly through a single module that implements common functionality. The general approach to combat entropy is refactoring operations, where code is rewritten to make it more modular and maintainable without changing its behaviour. The main risk of such operations is breaking existing functionality, in particular in a project the size of CoSMoMVPA (over 10,000 lines of code), where changes in one function may result in unexpected behavioural changes in another function. We use a test suite that can be run using either xUnit (Eddins, 2013; on the Matlab platform only) or MOxUnit (Oosterhof, 2015b; runs on both Matlab and GNU Octave). Through the use of a test suite with high (above 90%) code coverage, any broken functionality is likely to be detected rapidly. As in PyMVPA, the test suite can be run manually and is also run automatically through an online continuous integration testing service. (As of 2016, we use the travis system, which provides free testing services for open source projects). Every time when code is (proposed to be) included in the official branch (through a `github` *pull request*) or updated directly (through `git push`), the suite is run automatically by the travis system, and developers notified by email if any test failed. (Changes in) code coverage are reported through a customly written MOcov toolbox (Oosterhof, 2015a) and the coveralls.io service. Through such continuous integration testing, changes in the code can be made with relatively high confidence, because if anything would break it is likely to be detected quickly through failing tests.

### 4.3 Limitations

While CoSMoMVPA aims to be versatile and easy to use, it does not provide a graphical user interface. There are two reasons for this. First, the wide variety of options available in CoSMoMVPA means that writing the code for a user interface would take significant resources. Second, using scripts instead of a GUI has the advantage that analyses are sharable and reproducible.

CoSMoMVPA also does not support data preprocessing and has limited visualization abilities. This is because other packages for fMRI (SPM, AFNI, BrainVoyager, FSL) and M/EEG (FieldTrip and EEGLAB) already do this. Instead, CoSMoMVPA provides support for a wide variety of formats, so that preprocessed data from these packages can be imported, and MVPA results can be exported for visualization and further analyses.

## 5 Conclusion

In the present work, we presented CoSMoMVPA, an open source Matlab / GNU Octave toolbox for MVPA that runs on open source platforms. It supports all commonly-used MVPA techniques and a wide variety of imaging formats. It can easily be integrated into existing pipelines, using existing preprocessed data as input for MVPA, and exportomg results for visualization or further analyses with other packages. Through a unified approach to MVPA of fMRI and M/EEG, a generalized searchlight allows for data-driven localization of effects across space, time, and frequencies. State-of-the-art correction for multiple comparisons is provided to maximize sensitivity while controlling for type-1 errors.

CoSMoMVPA aims to provide basic building blocks for classification analysis, representational similarity analysis, the time generalization method, and generalized searchlight analyses. These building blocks can be combined to form sophisticated analysis pipelines. Its architecture is versatile, modular and flexible, which we believe makes it easy to use for non-experts, and extensible by more advanced users for purposes not envisioned by the developers.

CoSMoMVPA is designed to accommodate future growth, and uses software engineering best practices for development and quality control, including code version control, an automated test suite with high code coverage, continuous integration testing, and a build system for documentation. As new algorithms for analysing neuroimaging data are designed and implemented, we believe they can easily be added in the future.

We hope that through its simple and consistent design, CoSMoMVPA lowers the barriers to adoption of advanced multivariate techniques, and provides more cognitive neuroscientists easy access to a wider and more effective set of tools for analyzing brain data, which in turn increases the pace of new discoveries about brain function.

## 6 Acknowledgements

We thank Gunnar Blohm and Sara Fabri for inviting two of the authors (NNO and ACC) to organize a workshop at the 2013 Summer School in Computational Sensory-Motor Neuroscience (CoSMo), which formed the basis of CoSMoMVPA; Gunnar Blohm for providing us with the CoSMo logo; Michael Hanke and Yaroslav Halchenko for their work on PyMVPA, which inspired the semantics and data structure of datasets as well as the development process; Yaroslav Halchenko, Matteo Visconti di Oleggio Castello and Jens Scharzbach for code contributions; Jens Scharzbach for help with improving and adding exercises; Stephania Bracci for providing a dataset in BrainVoyager format; Nathan Weisz and Gianpaolo Demarchi for providing an M/EEG dataset; Jimmy Shen for providing a free/open source NIFTI library; Joern Diedrichsen and Tobias Wiestler for contributions to the LDA classifier and the free/open source surfing toolbox; Guillaume Flandin for providing the free/open source GIfTI library for Matlab; Robert Oostenveld and colleagues for providing the free/open source FieldTrip package; Ziad Saad and Gang Chen for providing the free/open source AFNI Matlab toolbox; Jochen Weber for providing the free/open source NeuroElf toolbox; Steve Eddins for providing the MATLAB xUnit Test Framework; and Thomas Smith for inspiration of the documentation testing functionality. We thank the following people for contributing valuable suggestions, advice, support, or code to CoSMoMVPA: Talia Brandman, Robert W. Cox, Sarah Belinda Aimee Degosciu, Hanna Gertz, Yaroslav Halchenko, Thomas Hartmann, Clayton Hickey, Daniel Kaiser, James Keidel, Sukhbinder Kumar, Cristina Lava, Seth Levine, Stefania Mattioni, Mike Miller, Sam Nastase, Nicholas Peatfield, Liuba Papeo, Alexis Perez, Jia Hou Poh, Daria Proklova, Reshanne Reader, Anne Roefs, Mohamed Tawfik, Luca Turella, and Moritz Wurm. We thank Karen Cuculiza, Scott Fairhall, and Marius Peelen for useful comments on an earlier draft of this manuscript. This work has been supported by the Autonomous Province of Trento, Italy, Call ‘Grandi Progetti 2012’, project ‘Characterizing and improving brain mechanisms of attention - ATTEND’.

### 7 Conflict of interest statement

The authors declare that the research was conducted in the absence of any commercial or financial relationships that could be construed as a potential conflict of interest.

## Footnotes

1 Throughout the manuscript we use the shorthand “M/EEG” to refer collectively to electrophysiological data including magneto-encephalography (MEG), scalp electro-encephalography (EEG), and intracranial EEG (iEEG), electro-corticography (ECoG).

